# Structural analyses of Trichomonas vaginalis pyrophosphate-dependent phosphofructokinase (TvPPi-PFK)

**DOI:** 10.64898/2026.03.28.715000

**Authors:** Avery Chiu, Lijun Liu, Steve Seibold, Kevin Battaile, Justin Craig, Elizabeth Harmon, Sandhya Subramanian, Lisabeth Cron, Bart Staker, Peter J Myler, Scott Lovell, Wesley C. Van Voorhis, Graham Chakafana, Oluwatoyin A. Asojo

## Abstract

*Trichomonas vaginalis* causes trichomoniasis, the most common non-viral sexually transmitted disease in humans. *T. vaginalis* pyrophosphate-dependent phosphofructokinase (*Tv*PPi-PFK) is a putative target for rational, structure-based drug discovery, given its absence in mammals and its importance for parasite survival. *Tv*PPi-PFK is a cytosolic enzyme that catalyzes the phosphorylation of fructose-6-phosphate using pyrophosphate (PPi) as the phosphoryl donor. This reversible reaction, catalyzed by *Tv*PPi-PFK, is the first committed step in glycolysis. Its reverse reaction is vital for gluconeogenesis in *T. vaginalis*. The purification, crystallization, structure determination, and crystal structures of *Tv*PPi-PFK are reported. *Tv*PPi-PFK is the first reported eukaryotic PPi-PFK structure. *Tv*PPi-PFK retains the overall PPi-PFK topology observed in bacterial PPi-PFK including conserved motifs essential for pyrophosphate binding and PPi-PFK catalytic activity. In addition to the catalytic PPi-PFK binding sites, *Tv*PPi-PFK has two additional ligand binding sites. The first binds AMP usurped during protein production and helps stabilize the *Tv*PPi-PFK tetramer. A second ligand binding site was observed in proximity to the AMP-binding site and accommodates sugar phosphates soaked into preformed crystals. This sugar phosphates binding site is distinct from the *Tv*PPi-PFK active site that binds fructose-6-phosphate. Future mutagenesis and activity studies are planned to determine the relevance of both sites.

**Synopsis:** The production, crystallization, and crystal structures of a pyrophosphate-dependent phosphofructokinase from *Trichomonas vaginalis* (*Tv*PPi-PFK) are reported. *Tv*PPi-PFK has a prototypical PPi-PFK active site as well as unexpected AMP and sugar-phosphate binding sites at the dimer interface.

## 1. Introduction

The unicellular parasitic protozoan *Trichomonas vaginalis* (*T. vaginalis*) causes ∼156 million new annual cases of trichomoniasis, the most prevalent non-viral sexually transmitted disease globally (Edwards *et al*., 2016, Satterwhite *et al*., 2013, Molgora *et al*., 2023). Approximately 3.7 million people in the USA have trichomoniasis, exceeding both chlamydia and gonorrhea combined and population studies show that the highest incidence is among incarcerated women (Satterwhite *et al*., 2013). *T. vaginalis* exclusively infects humans, and trichomoniasis leads to higher rates of HIV, infertility, pre-term birth, HPV, cervical, and prostate cancer (Tsang *et al*., 2019, Van Gerwen & Muzny, 2019, Zhang *et al*., 2022). *T. vaginalis* is anaerobic, thriving in the urethra, vagina, and vulva, and is readily spread by asymptomatic people with ∼1.1 million new infections annually in the USA (Flagg *et al*., 2019, Meites, 2013, Van Gerwen *et al*., 2023). Trichomoniasis is most often treated with oral antibiotics: a single dose of 2 g of tinidazole, a 7-day course of twice-daily 500mg metronidazole, or a single or 5-day course of secnidazole (Muzny *et al*., 2022). Drug resistance to metronidazole was identified over 60 years ago (Robinson, 1962). Current treatments often fail if alcohol is consumed, and reinfection is common if partners are not treated, and allergic reactions may affect compliance (Muzny *et al*., 2022). There is therefore a need to identify alternative new therapeutics and therapeutic targets for trichomoniasis.

*T. vaginalis* is a priority infectious disease for structural studies by the Seattle Structural Genomics Center for Infectious Disease (SSGCID) because of its clinical significance. SSGCID studies include the determination of the structures of potential new *T. vaginalis* therapeutic targets and vital metabolic enzymes such as pyrophosphate-fructose 6-phosphate 1-phosphotransferase 1 (PFP-1). PFP-1, also referred to as pyrophosphate-dependent phosphofructokinase (PPi-PFK), catalyzes the first committing step in glycolysis, the phosphorylation of fructose 6-phosphate. PPi-PFK uses pyrophosphate (PPi) as the phosphoryl donor to catalyze the synthesis of fructose 1,6-bisphosphate and inorganic phosphate. Since the reaction is reversible, PPi-PFK functions in both glycolysis and gluconeogenesis. This is a deviation from the otherwise well-known reaction in glycolysis catalyzed by ATP-PFK in which adenosine triphosphate (ATP) is used as the phosphoryl donor instead of PPi (Scheme 1) (Steinbuchel & Muller, 1986, Schneider *et al*., 2011, Rada *et al*., 2015). A review of the diversity and complexity of the structure-function relationships of members of the PFK superfamily was published in 2025 (Compton & Patrick, 2025).

*T. vaginalis* has both PPi-PFK and ATP-PFK enzymes (Steinbuchel & Muller, 1986, Schneider *et al*., 2011, Rada *et al*., 2015). *Tv*PPi-PFK is a cytosolic enzyme responsible for most of *T. vaginalis’* glycolytic flux and energetic needs (Rada *et al*., 2015). *T. vaginalis* has four ATP-PFK enzymes, which have less than 2% of the activity of the PPi-PFK (Rada *et al*., 2015, Schneider *et al*., 2011). Furthermore, all four *Tv*ATP-PFK are localized in the hydrogenosomes, which are double membrane organelles that *T. vaginalis* possesses instead of mitochondria (Steinbuchel & Muller, 1986, Schneider *et al*., 2011, Rada *et al*., 2015). Full lenght *Tv*PPi-PFK is a 426 amino acid protein of ∼48kDa MW. *Tv*PPi-PFK shares less than 29% sequence identity with any reported structure in the protein data bank. A blast search reveals *Tv*PPi-PFK has no appreciable sequence similarity to any human proteins. Due to its importance for *T. vaginalis* survival and lack of similarity to human proteins, *Tv*PPi-PFK is a target for rational drug discovery by the SSGCID. As part of efforts to understand this putative drug target, we present the purification, crystallization, and crystal structure of *Tv*PPi-PFK.

## 2. Materials and methods

### 2.1. Macromolecule production

*Tv*PPi-PFK was cloned, expressed, and purified using standard methods developed by the SSGCID (Bryan *et al*., 2011, Choi *et al*., 2011, Serbzhinskiy *et al*., 2015, Makori *et al*., 2026, Nair *et al*., 2026, Srivastava *et al*., 2026, Austin *et al*., 2026). The full-length gene for pyrophosphate-fructose 6-phosphate 1-phosphotransferase 1 (PFP-1) from *Trichomonas vaginalis ATCC PRA-98 / G3* (Uniprot A2DXT4) encoding amino acids 1-426 was PCR-amplified from gDNA using the primers shown in Table 1, cloned into the BG1861 and then the plasmid DNA was transformed into chemically competent *Escherichia coli* BL21(DE3) cells. After testing for expression, 2L of culture was grown using auto-induction media (Studier, 2005) in a LEX Bioreactor (Epiphyte Three) as described previously (Serbzhinskiy *et al*., 2015). N-terminal hexa-histidine tagged *Tv*PPi-PFK was purified using the standardized previously described two-step protocol of an immobilized metal (Ni^2+^) affinity chromatography (IMAC) step followed by a preparative size-exclusion chromatography (SEC) on an AKTApurifier 10 (GE Healthcare) using automated IMAC and SEC programs (Serbzhinskiy *et al*., 2015). Briefly, thawed bacterial pellets (25 g) were lysed by sonication in 200 ml lysis buffer (25 mM HEPES pH 7.0, 500 mM NaCl, 5% (v/v) glycerol, 0.5% (w/v) CHAPS, 30 mM imidazole, 10 mM MgCl_2_, 400 mg/ml AEBSF and 0.025% (w/v) sodium azide. After sonication, the crude lysate was incubated with 20 ml of benzonase (25 units/ml) at room temperature for 45 min with mixing. After clarifying the lysate by centrifugation (∼10,000 g for 1 hour using a Sorvall centrifuge, Thermo Scientific), the clarified supernatant was passed over a Ni-NTA HisTrap FF 5 ml column (GE Healthcare), which was pre-equilibrated with wash buffer (25 mM HEPES pH 7.0, 500 mM NaCl, 5% (v/v) glycerol, 30 mM imidazole). The column was washed with 20 column volumes (CV) of wash buffer and eluted with elution buffer (20 mM HEPES, pH 7.0, 500 mM NaCl, 5% (v/v) glycerol, 500 mM imidazole) over a 7CV linear gradient. Peak fractions were pooled, concentrated to ∼5 ml and loaded onto a Superdex 75 26/60 column (GE Biosciences) attached to an ÄKTA Prime-plus FPLC system (GE Biosciences) that was equilibrated with storage buffer (20 mM HEPES, pH 7.0, 300 mM NaCl, 5% glycerol, and 1 mM TCEP). The peak fractions were collected and assessed for purity by SDS-PAGE. *Tv*PPi-PFK eluted as a broad peak, accounting for ∼90% of the protein product. The purest fractions had molecular mass ∼48 kDa close to the theoretical molecular weight 47.6 kDa of recombinant Histagged *Tv*PPi-PFK as assessed by SDS page gel using molecular weight standards (Figure S.1). These fractions were pooled and concentrated to ∼25 mg/ml with an Amicon purification system (Millipore). 110 *µ*l aliquots of recombinant *Tv*PPi-PFK were flash-frozen in liquid nitrogen and stored at -80°C until used.

**Table 1.**
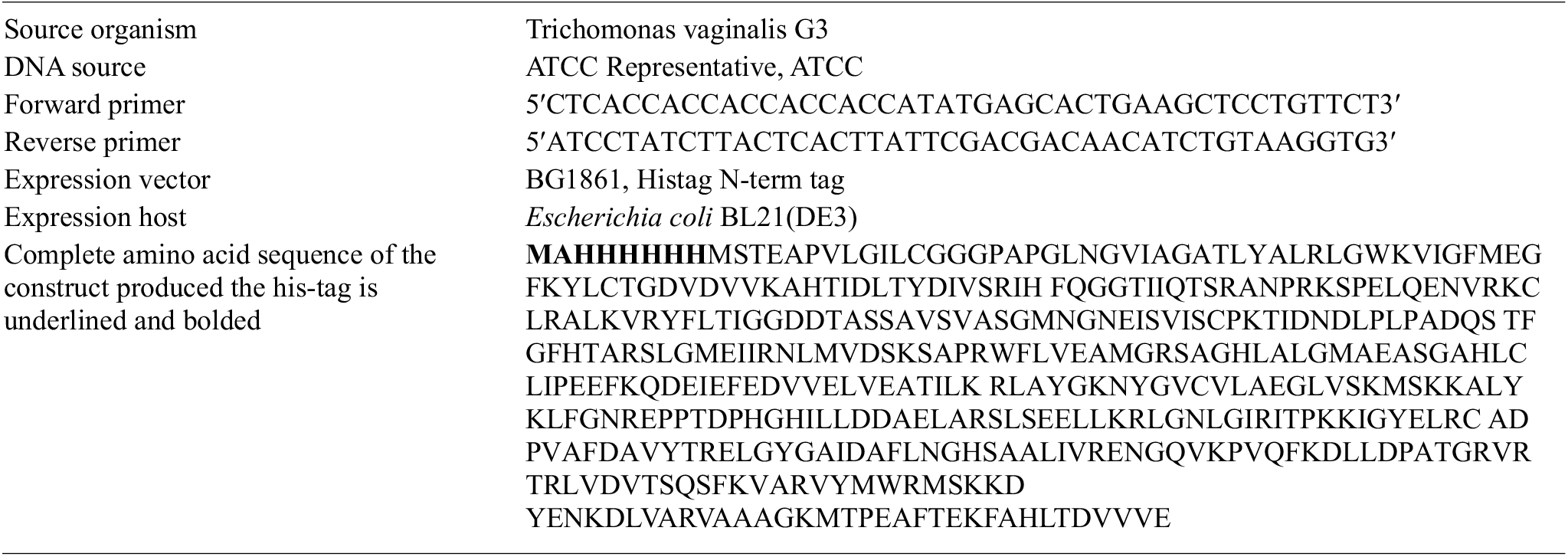
Macromolecule production information.

### 2.2. Crystallization

All crystallization experiments were conducted using an NT8 drop-setting robot (Formulatrix Inc.) and UVXPO MRC (Molecular Dimensions) sitting-drop vapor diffusion plates at 17 ^°^C. *Tv*PPi-PFK crystals grew directly from the Molecular Dimensions crystallization screen condition Morpheus A4 (Gorrec, 2009). Briefly, 200 nL of protein in storage buffer and 200 nL of crystallization solution (Morpheus A4: 12.5% (v/v) MPD, 12.5% (v/v) PEG 1000, 12.5% (w/v) PEG 3350, 100 mM Imidazole/MES, pH 6.5, 30 mM MgCl_2_ and 30 mM CaCl_2_) were added to the sitting drop and equilibrated against reservoir containing 40 mL of crystallization solution (Table 2). The crystal was soaked in soaking solution (5 mM ATP and 5 mM pyrophosphate (PPi), 12.5% (v/v) MPD, 12.5%(v/v) PEG 1000, 12.5% (w/v) PEG 3350, 100 mM Imidazole/MES, pH 6.5, 30 mM MgCl_2_ and 30 mM CaCl_2_) overnight (at least 8 hours). The crystal was vitrification in liquid Nitrogen, mouted on pucks and shipped to Brookhaven National Laboratory for remote data collection.

**Table 2.** Crystallization and soaking conditions.

|  |  |  |  |  |
| --- | --- | --- | --- | --- |
| 235 | **Accession code** | 9MDT (pdb_00009mdt) | 9MF6 (pdb_00009mf6) | 9MED (pdb_00009med) |
| 236 | **Method** | Vapor diffusion, sitting drop | Vapor diffusion, sitting drop | Vapor diffusion, sitting drop |
| 237 | **Plate type** | Plate Liu-S-143 AB34 | Plate Liu-S-143 AB34 | Plate Liu-S-143 AB34 |
| 238 | **Temperature (K)** | 291 | 291 | 291 |
| 239 | **Protein concentration** | 25 mg mL<sup>-1</sup> | 25 mg mL<sup>-1</sup> | 25 mg mL<sup>-1</sup> |
| 240 | **Buffer composition of protein solution** | 20 mM HEPES, pH 7.0, 300 mM NaCl, 5% glycerol, and 1 mM TCEP | 20 mM HEPES, pH 7.0, 300 mM NaCl, 5% glycerol, and 1 mM TCEP | 20 mM HEPES, pH 7.0, 300 mM NaCl, 5% glycerol, and 1 mM TCEP |
| 241 | **Composition of reservoir solution** | Morpheus A4: 12.5%(v/v) MPD, 12.5%(v/v) PEG 1000, 12.5% (w/v) PEG 3350, 100 mM Imidazole/MES, pH 6.5, 30 mM MgCl<sub>2</sub> and 30 mM CaCl<sub>2</sub> | Morpheus A4: 12.5%(v/v) MPD, 12.5%(v/v) PEG 1000, 12.5% (w/v) PEG 3350, 100 mM Imidazole/MES, pH 6.5, 30 mM MgCl<sub>2</sub> and 30 mM CaCl<sub>2</sub> | Morpheus A4: 12.5%(v/v) MPD, 12.5%(v/v) PEG 1000, 12.5% (w/v) PEG 3350, 100 mM Imidazole/MES, pH 6.5, 30 mM MgCl<sub>2</sub> and 30 mM CaCl<sub>2</sub> |
| 242 | **Volume and ratio of drop** | 0.2 μl : 0.2 μl | 0.2 μl : 0.2 μl | 0.2 μl : 0.2 μl |
| 243 | **Volume of reservoir** | 40 μl | 40 μl | 40 μl |
| 244 | **Soaking of crystal in solution containing** | 5 mM ATP and 5 mM pyrophosphate (PPi), 12.5%(v/v) MPD, 12.5%(v/v) PEG 1000, 12.5% (w/v) PEG 3350, 100 mM Imidazole/MES, pH 6.5, 30 mM MgCl<sub>2</sub> and 30 mM CaCl<sub>2</sub> | 2 mM ADP and a-D-Ribose 5-phosphate (R5P), 12.5%(v/v) MPD, 12.5%(v/v) PEG 1000, 12.5% (w/v) PEG 3350, 100 mM Imidazole/MES, pH 6.5, 30 mM MgCl<sub>2</sub> and 30 mM CaCl<sub>2</sub> | 2 mM AMP, b-D-Glucose-6-phosphate (G6P), 12.5%(v/v) MPD, 12.5%(v/v) PEG 1000, 12.5% (w/v) PEG 3350, 100 mM Imidazole/MES, pH 6.5, 30 mM MgCl<sub>2</sub> and 30 mM CaCl<sub>2</sub> |

### 2.3. Data collection, processing, structure determination and refinement

Diffraction data were collected at 100 K on a DECTRIS EIGER2 XE 9M detector at NSLS-II BEAMLINE 19-ID at Brookhaven National Laboratory. Data were integrated with XDS and reduced with XSCALE (Kabsch, 2010). Raw X-ray diffraction images are stored at the Integrated Resource for Reproducibility in Macromolecular Crystallography at https://www.proteindiffraction.org. The structures were determined by molecular replacement with Phaser (McCoy *et al*., 2007) from the CCP4 suite of programs (Collaborative Computational Project, 1994, Krissinel *et al*., 2004, Winn *et al*., 2011, Agirre *et al*., 2023). The molecular replacement search model pdb entry 9DQM. The ligands were modeled The structures were refined by iterative model building with COOT (Emsley *et al*., 2010) and PHENIX (Adams *et al*., 2010, Liebschner *et al*., 2019). Structure quality was checked with MolProbity (Williams *et al*., 2018). Data-reduction and refinement statistics are shown in Table 3. Coordinates and structure factors is deposited with the Worldwide PDB (wwPDB).

**Table 3.**
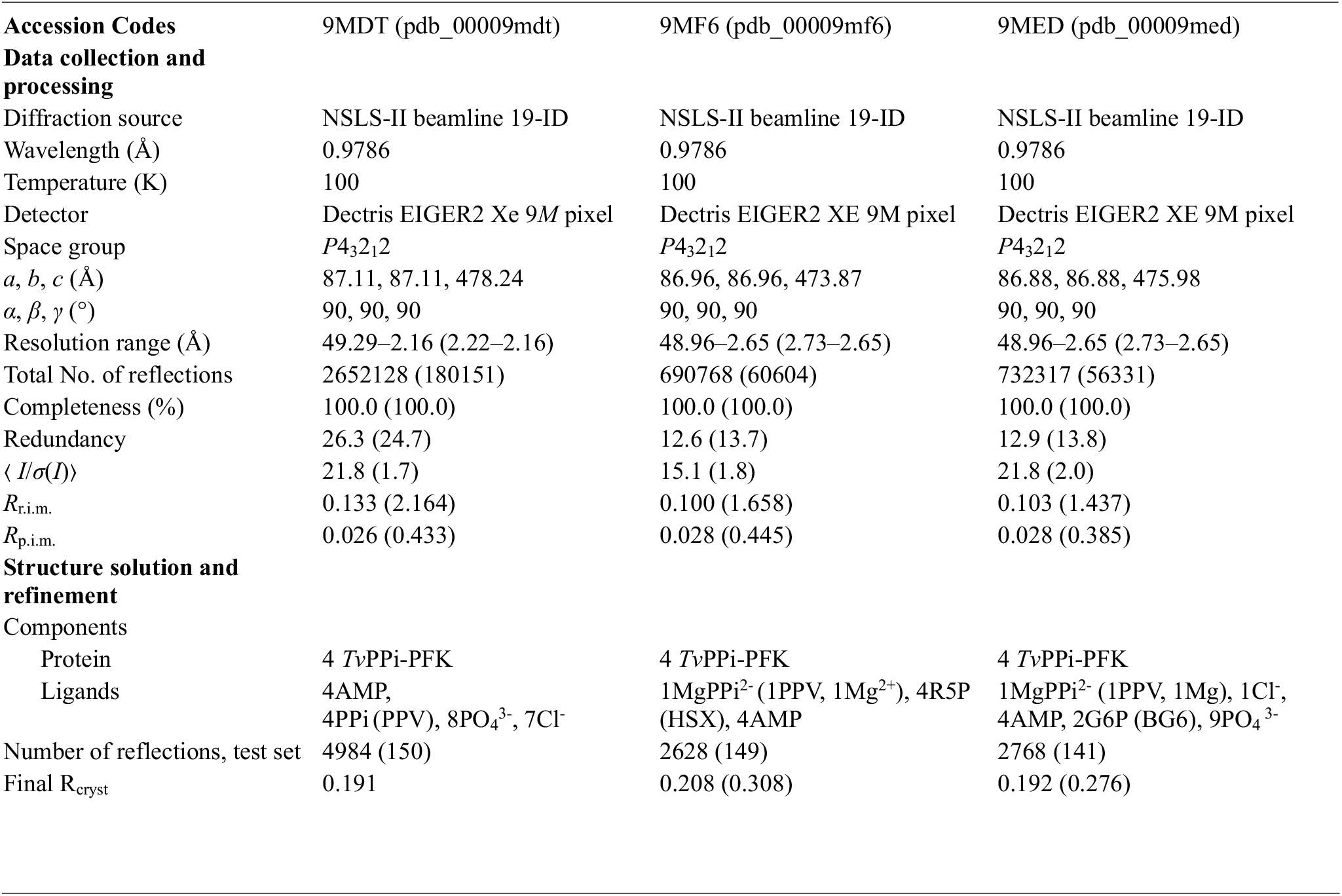
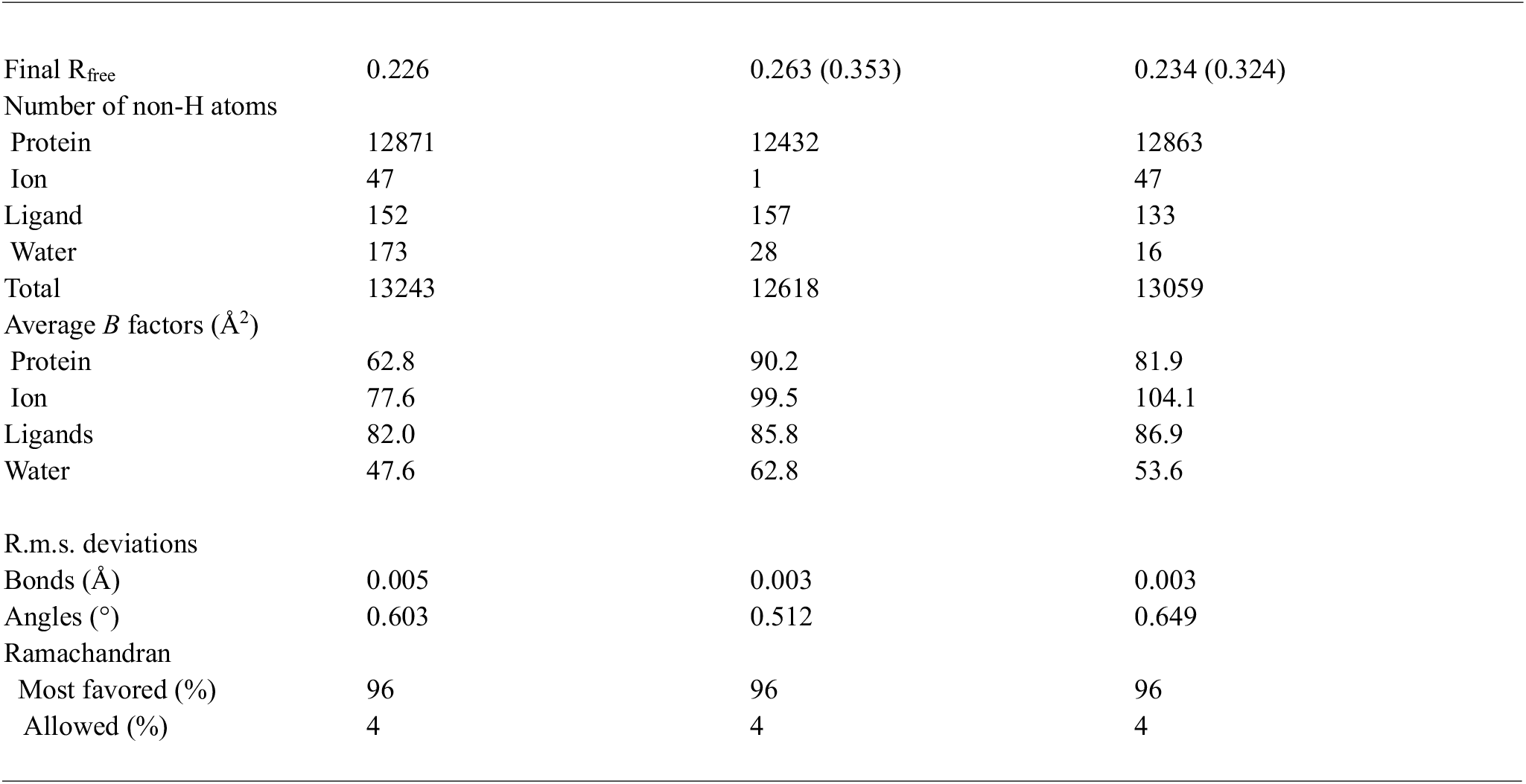
Data collection, processing and refinement (Values for the outer shell are given in parentheses).

|  |  |  |  |  |
| --- | --- | --- | --- | --- |
| 247 | **Accession Codes** | 9MDT (pdb_00009mdt) | 9MF6 (pdb_00009mf6) | 9MED (pdb_00009med) |
| 248 | **Data collection and processing** |  |  |  |
| 249 | **Diffraction source** | NSLS-II beamline 19-ID | NSLS-II beamline 19-ID | NSLS-II beamline 19-ID |
| 250 | **Wavelength (Å)** | 0.9786 | 0.9786 | 0.9786 |
| 251 | **Temperature (K)** | 100 | 100 | 100 |
| 252 | **Detector** | Dectris EIGER2 Xe 9M pixel | Dectris EIGER2 XE 9M pixel | Dectris EIGER2 XE 9M pixel |
| 253 | **Space group** | *P*<sub>4</sub><sub>3</sub><sub>2</sub><sub>1</sub><sup>2</sup> | *P*<sub>4</sub><sub>3</sub><sub>2</sub><sub>1</sub><sup>2</sup> | *P*<sub>4</sub><sub>3</sub><sub>2</sub><sub>1</sub><sup>2</sup> |
| 254 | ***a*, *b*, *c* (Å)** | 87.11, 87.11, 478.24 | 86.96, 86.96, 473.87 | 86.88, 86.88, 475.98 |
| 255 | ***α*, *β*, *γ* (°)** | 90, 90, 90 | 90, 90, 90 | 90, 90, 90 |
| 256 | **Resolution range (Å)** | 49.29–2.16 (2.22–2.16) | 48.96–2.65 (2.73–2.65) | 48.96–2.65 (2.73–2.65) |
| 257 | **Total No. of reflections** | 2652128 (180151) | 690768 (60604) | 732317 (56331) |
| 258 | **Completeness (%)** | 100.0 (100.0) | 100.0 (100.0) | 100.0 (100.0) |
| 259 | **Redundancy** | 26.3 (24.7) | 12.6 (13.7) | 12.9 (13.8) |
| 260 | ***⟨I/σ(I)⟩*** | 21.8 (1.7) | 15.1 (1.8) | 21.8 (2.0) |
| 261 | ***R*<sub>r.i.m.</sub>** | 0.133 (2.164) | 0.100 (1.658) | 0.103 (1.437) |
| 262 | ***R*<sub>p.i.m.</sub>** | 0.026 (0.433) | 0.028 (0.445) | 0.028 (0.385) |
| 263 | **Structure solution and refinement** |  |  |  |
| 264 | **Components** |  |  |  |
| 265 | Protein | 4 TvPPi-PFK | 4 TvPPi-PFK | 4 TvPPi-PFK |
| 266 | Ligands | 4AMP, 4PPi (PPV), 8PO<sub>4</sub><sup>3-</sup>, 7Cl<sup>-</sup> | 1MgPPi<sup>2-</sup> (1PPV, 1Mg<sup>2+</sup>), 4R5P (HSX), 4AMP | 1MgPPi<sup>2-</sup> (1PPV, 1Mg), 1Cl<sup>-</sup>, 4AMP, 2G6P (BG6), 9PO<sub>4</sub><sup>3-</sup> |
| 267 | **Number of reflections, test set** | 4984 (150) | 2628 (149) | 2768 (141) |
| 268 | **Final *R*<sub>cryst</sub>** | 0.191 | 0.208 (0.308) | 0.192 (0.276) |
| 269 | Final $R_{\text{free}}$ | 0.226 | 0.263 (0.353) | 0.234 (0.324) |
| 270 | Number of non-H atoms |  |  |  |
| 271 | Protein | 12871 | 12432 | 12863 |
| 272 | Ion | 47 | 1 | 47 |
| 273 | Ligand | 152 | 157 | 133 |
| 274 | Water | 173 | 28 | 16 |
| 275 | Total | 13243 | 12618 | 13059 |
| 276 | Average $B$ factors ($\text{\AA}^2$) | | | |
| 277 | Protein | 62.8 | 90.2 | 81.9 |
| 278 | Ion | 77.6 | 99.5 | 104.1 |
| 279 | Ligands | 82.0 | 85.8 | 86.9 |
| 280 | Water | 47.6 | 62.8 | 53.6 |
| 281 |  |  |  |  |
| 282 | R.m.s. deviations |  |  |  |
| 283 | Bonds ($\text{\AA}$) | 0.005 | 0.003 | 0.003 |
| 284 | Angles ($^\circ$) | 0.603 | 0.512 | 0.649 |
| 285 | Ramachandran |  |  |  |
| 286 | Most favored (%) | 96 | 96 | 96 |
| 287 | Allowed (%) | 4 | 4 | 4 |

**Table 4.** Scheme 1: Comparison of ATP-PFK and PPi-PFKs.

|  |  |  |  |
| --- | --- | --- | --- |
| 290 | **Enzyme** | **ATP-PFK (EC 2.7.1.11)** | **PPi-PFK (EC 2.7.1.90)** |
| 291 | **Phosphate source** | ATP | PPi |
| 292 | **Reaction** | $\text{ATP} + \text{F6P} \rightarrow \text{ADP} + \text{F1,6BP}$ | $\text{PPi} + \text{F6P} \rightleftharpoons \text{Pi} + \text{F1,6BP}$ |
| 293 | **Thermodynamics** | Irreversible in vivo | Near-equilibrium, reversible |
| 294 | **Regulation** | Major glycolytic regulatory point | PPi homeostasis, glycolysis, gluconeogenesis, etc |
| 295 | **Typical organisms that utilize** | Animals, fungi, many bacteria | Plants, protists, archaea, anaerobic parasites |

### 2.4. Mass Photometry

Mass photometry measures the molecular weight and relative abundance of proteins in a sample with light scattering for an analytical range of 40 kDa - 5 MDa (Wu & Piszczek, 2021, Young *et al*., 2018). Standard methods established by Dr. Lisa Tuttle (University of Washington) were followed. 24×50 mm glass coverslips were cleaned with isopropanol and dried with nitrogen gas. The TwoMP Mass Photometer instrument (Refeyen Ltd, Oxford, UK) was calibrated with 50 nM b-amylase. 10 μl of storage buffer (20 mM HEPES, 300 mM NaCl, 5% (v/v) glycerol, and 1 mM TCEP) was measured in sample gasket wells, followed by 10 μl of the ∼50 nM protein samples at Room Temperature. The first condition was recombinant ∼50 nM *Tv*PPi-PFK in storage buffer. The second condition is ∼50 nM recombinant *Tv*PPi-PFK in 5 mM ATP, 5 mM PPi, 5 mM MgCl_2_, 20 mM HEPES, 300 mM NaCl, 5% (v/v) glycerol, and 1 mM TCEP. Mass histograms were collected with the DiscoverMP analysis software and analyzed with single Gaussian fits in IGOR-Pro v.8 (Wavemetrics, Lake Oswego, OR).

## 3. Results and discussion

### Overall structure

The crystallized *Tv*PPi-PFK protein included 8 additional vector-derived N-terminus amino acid residues, including a hexahistidine tag (Table 1). These vector-derived N-terminal residues are disordered and were not modeled. Most mature polypeptide main-chain and side-chain residues were ordered and built and the refined structures of *Tv*PPi-PFK contains a homotetramer in the asymmetric unit (Table 3). The *Tv*PPi-PFK structures are modelled as complexes with inorganic pyrophosphate (PPi), adenosine monophosphate (AMP), phosphate (PO_4_ ^3-^), a-D-Ribose 5-phosphate (R5P), or *β*-D-glucose 6-phosphate (G6P), magnesium, and chloride ions (Table 3). Recombinant *Tv*PPi-PFK is expressed in media that contains phosphate, magnesium, and chloride ions (Studier, 2005). The crystallization solution also contains magnesium and chloride ions. No exogenous PPi or AMP was added to the purification or crystallization solutions prior to the soaking experiments. It is likely that the tetramers that crystallized are those with AMP bound since crystallization is a purification step. Soaking with ATP or ADP or inhibitors could not displace the bound AMP, thus all the crystal structures are complexes with AMP.

Each polypeptide chain of *Tv*PPi-PFK adopts the prototypical topology of a PPi-PFK, with the pyrophosphate-binding site sitting between the N-terminal and C-terminal domains (Figure 1a). PROMOTIF analysis reveals that each *Tv*PPi-PFK protomer’s secondary structure is ∼15% strand, ∼41% alpha helix, 7% 3-10 helix. Each *Tv*PPi-PFK chain assembles into 2 beta-sheets (a 7-stranded mixed sheet and a 4-stranded parallel sheet). 3 beta-alpha-beta motifs, and 2 beta hairpins. The asymmetric unit contains four copies of the polypeptide (chains A-D), forming a dimer of two tightly packed dimers (Figures 1b - g). All the polypeptide chains are very similar, with an rmsd of ∼0.2 Å when superimposing all main-chain atoms. The asymmetric unit contains a homotetramer with an estimated molecular weight of ∼193kDa, suggestive of the canonical tetrameric PPi-PFK fold. The homotetramer has four large dimer interfaces (Figure 1). The larger dimer between chains A and chain D (equivalent to chains C and B) has a buried surface area of ∼2230 Å^2^. This is approximately double the ∼1270 Å^2^ buried surface area of the smaller dimer interface between chains A and chain C (equivalent to chains D and B).

**Figure 1.**
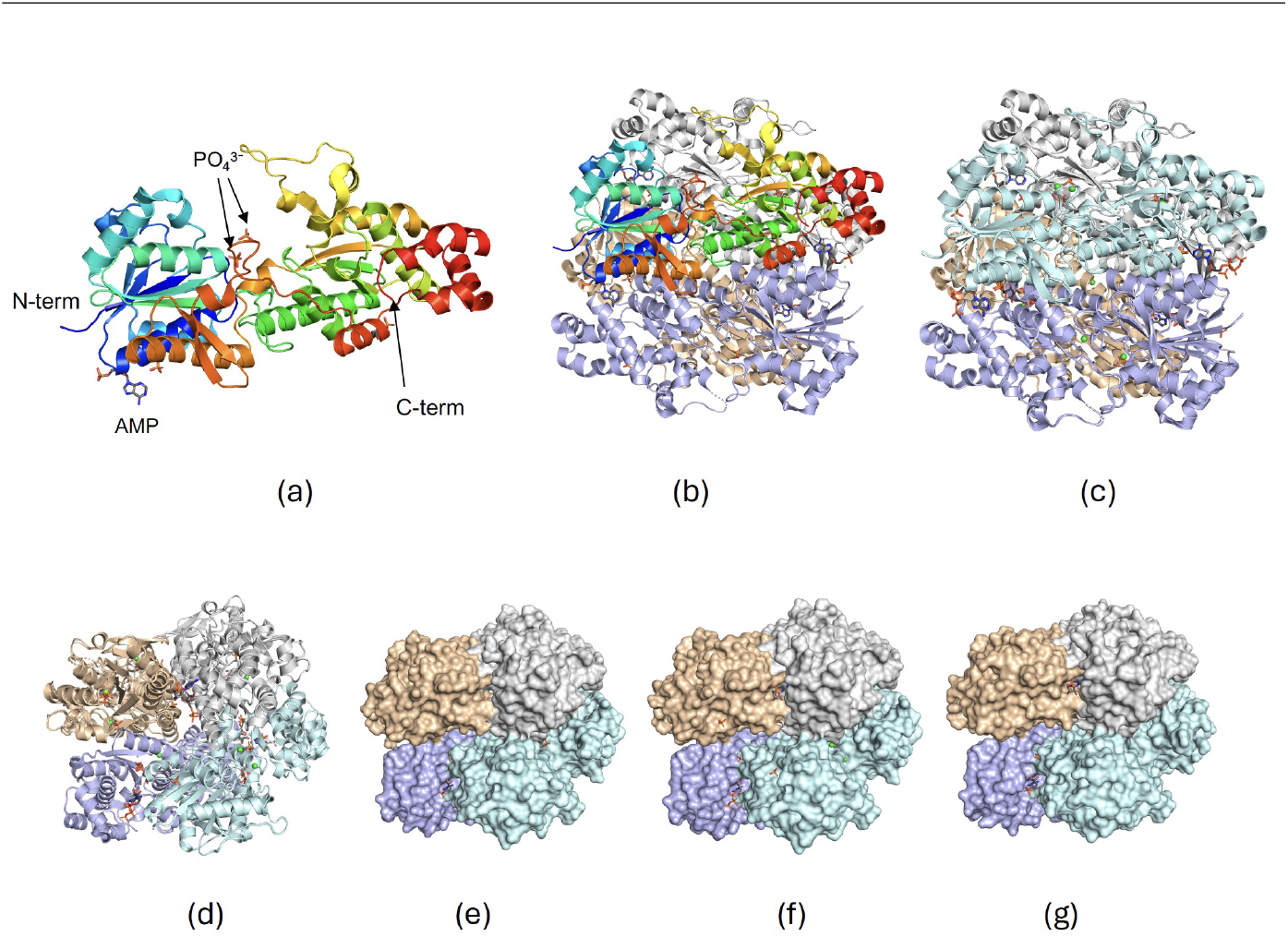
Overall structure of *Tv*PPi-PFK. (a) A representative protomer of *Tv*PPi-PFK (pdb entry 9MED, chain B, colored in rainbow from blue N-to red C-term) reveals the prototypical two-domain topology of PPi-PFK. Two phosphate molecules sit in the central substrate-binding cavity. Other ligands in the protomer are shown in sticks or spheres. (b) A representative *Tv*PPi-PFK homotetramer with one monomer is shown as a rainbow cartoon (same orientation as figure 1a) while the other three are shown in gray, tan, and blue. (c) Superposed tetramers *Tv*PPi-PFK (pdb entries 9MED, and 9MF6) reveal extensive structural similarity, chain B shown in cyan, chain C in gray, chain A in wheat, and chain D in blue. (d) An alternative view of the superposed tetramers reveals prototypical PPi-PFK homotetramer topology. The homotetramer surfaces plots of *Tv*PPi-PFK reveal large dimer interfaces for (e) 9MED and (f) 9MF6; some of the bound ligands in the dimer interface are visible from this orientation.

*Tv*PPi-PFK is predicted to be a biological tetramer based on Proteins, Interfaces, Structures, and Assemblies (PDBePISA) analysis (Krissinel, 2015). Mass photometry data experiments in the storage buffer reveals that recombinant *Tv*PPi-PFK is mainly tetrameric with smaller monomer and dimer peaks (Figure 2a). The addition of ATP, MgCl_2_, and pyrophosphate (PPi) reduces the monomer and dimer peaks to almost neglible level (Figure 2b). ATP stabilizes the tetramer even in the presence of EDTA (Figure 2c). Addition of ATP is required to reduce the relative aboundance of monomer and dimers and increase that of tetramers in solution (Figures 2a - d).

**Figure 2.**
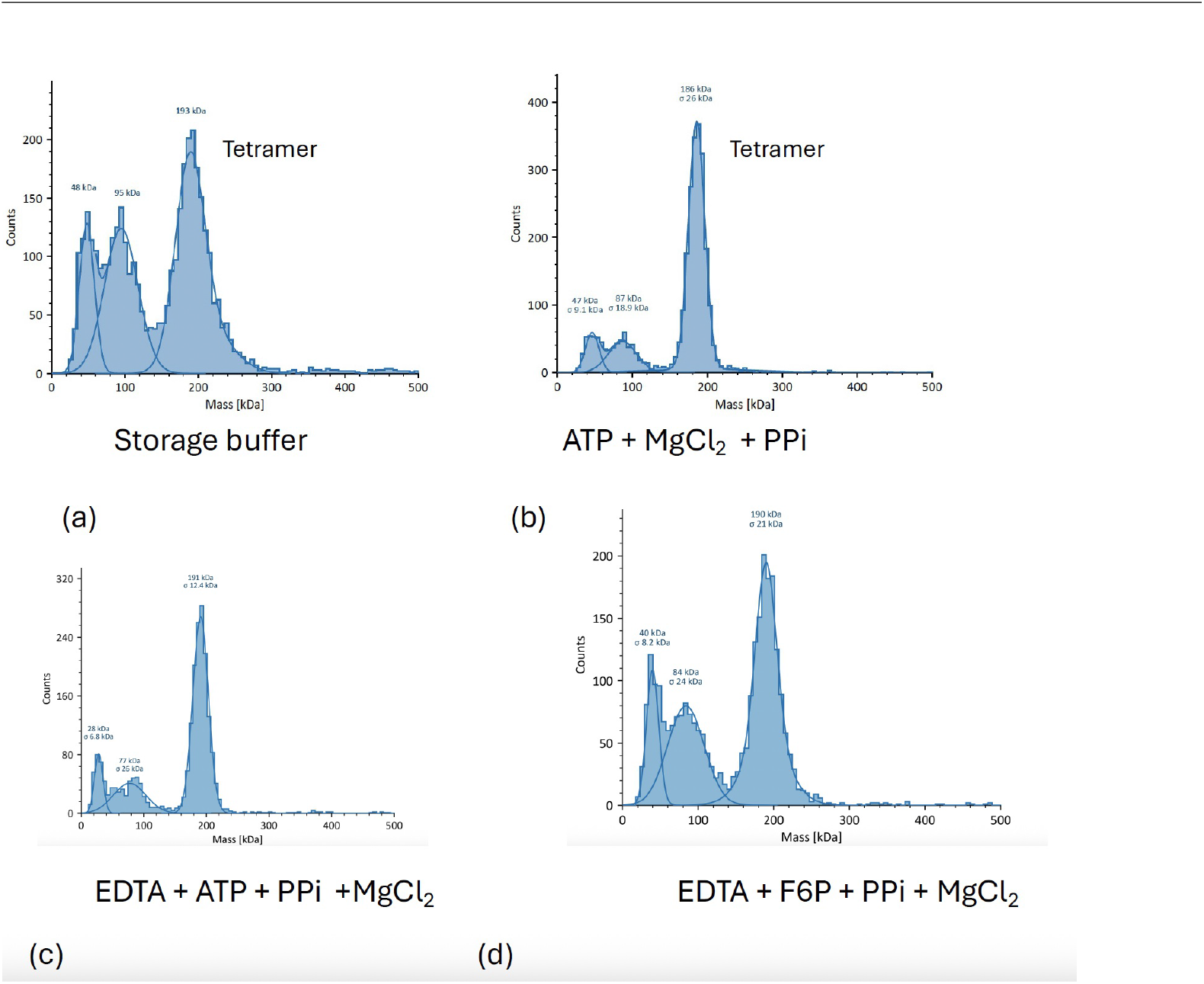
Mass Photometry analysis of recombinant *Tv*PPi-PFK. (a) Recombinant *Tv*PPi-PFK forms a mixture of tetramers, dimers, and monomers in storage buffer. (b) *Tv*PPi-PFK is mainly tetramer in buffer containing 5mM ATP, 5mM PPi and 5mM MgCl_2_ . (c) *Tv*PPi-PFK is mostly tetrameric in buffer containing 5mM EDTA, 5mM ATP, 5mM PPi and 5mM MgCl_2._ (e) recombinant *Tv*PPi-PFK is a mixture of tetramers, dimers, and monomers in buffer containing 5mM EDTA, 5mM F6P, 5mM PPi and 5mM MgCl_2._

### PPi binding and PPi-PFK motifs

The *Tv*PPi-PFK structures validate *Tv*PPi-PFK’s previously described PPi-PFK enzymatic activity (Mertens *et al*., 1989). PPi binds in the substrate-binding cavity and forms close interactions with Gly14, Gly15, Arg83, Gly113, Thr116, Asp115, and Lys139 (Figure 3a-d, Figure S.2). The residues involved in PPi binding are key classical pyrophosphate-dependent phosphofructokinase residues and motifs. *Tv*PPi-PFK has the canonical PPi-PFK pyrophosphate binding glycine (G15). The substrate-binding site contains the prototypical GGDD motif that ensures PPi specificity. The GGDD motif includes D114, which is important for Mg^2+^ binding, and D115, which is important for catalytic activity and stabilizes the transition state when PPi is the phosphoryl donor. A second key conserved PPi-PFK motif is the “KTID” substrate binding site-motif, which contains the catalytic lysine (Lys139). Lys139 is positioned to stabilize the transition state when PPi is the phosphoryl donor, while Asp142 serves as a proton donor (Figure S.2, Figure 3).

**Figure 3.**
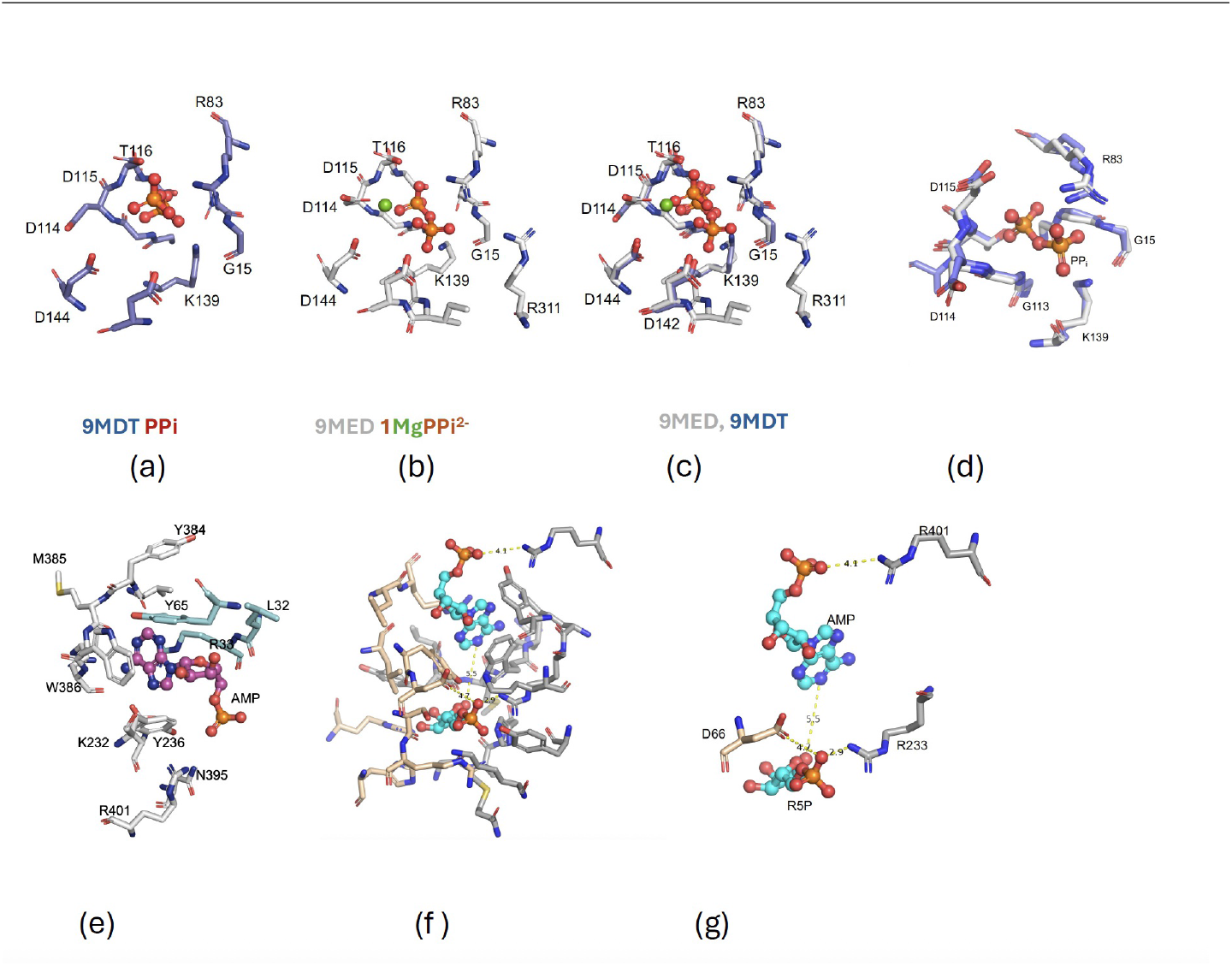
Ligand-binding by *Tv*PPi-PFK. Comparison of PPi binding by *Tv*PPi-PFK in the (a) absence and (b) presence of magnesium (c) superpositioning of the states. (d) Comparison of *Tv*PPi-PFK’s PPi-binding site with PPi (chain C of pdb entry 9MDT) and without PPi (Chain C of pdb entry MED). (e) ATP binding by *Tv*PPi-PFK is stabilized by residues from the protomer (gray) and from across the dimer interface (green). (f) Alternative view of *Tv*PPi-PFK’s putative AMP-binding site (pdb entry 4MF6) showing proximity of R5P and AMP and network of amino acid residues within 5 Å. (g) same view showing fewer residues.

### AMP and sugar phosphate binding

Recombinant *Tv*PPi-PFK is expressed in media that contains phosphate, magnesium, and chloride ions (Studier, 2005). The crystallization solution also contains magnesium and chloride ions. No exogenous PPi, ATP, or AMP was added in purification, or crystallization solutions prior to the soaking experiments. When *Tv*PPi-PFK crystals are soaked with only 2mM pyrophosphate, the resulting crystal structure pdb entry 9DQM | pdb_00009dqm contains AMP and PPi, suggesting that AMP is usurped during protein production. AMP molecules sit at the dimer interfaces and interact with the amino acid residues Leu32, Arg33, Tyr65 from one monomer and Lys232, Tyr236, Met385, Trp386 from a second monomer across the dimer interface (Figure 3a). Trp 386 forms a pi-stacking interaction with the adenine group, while the other residues form pi-stacking, hydrogen bonding, and other interactions with the ATP or AMP (Figures 3a, b). AMP binding at *Tv*PPi-PFK’s dimer interface suggests that it stabilizes the crystallographic dimer and we plan to conduct mutagenesis as well as enzymatic activity studies to determine if AMP/ADP/ATP affects *Tv*PPi-PFK’s PPi-PFK activity. Soaking preformed *Tv*PPi-PFK crystals with different sugar phosphates serendipitously resulted in the identification of a sugar phosphate binding site less than 5Å from the AMP-binding site (Figure 3e-f). These sugar phosphate sites are distinct from *Tv*PPi-PFK binding cavity which binds The 9MF6 structure contains four R5P molecules in the four newly identified sugar phosphate sites. Whereas the 9MED structure contains two G6P molecules occupying two of the four equivalent interface sites. LIGPLOT+ analysis indicates all R5P and G6P molecules form similar interactions with *Tv*PPi-PFK (Figure S.3). In structures lacking sugar phosphates (9MDT), phosphate ions occupy positions corresponding to the phosphoryl groups of the sugar phosphates (9MF6). The R5P/G6P binding-site is distinct from the PPi-PFK catalytic site that binds fructose 6-phosphate (Figure S.3).

### Comparison to structural neighbors

ENDScript (Gouet *et al*., 2003, Robert & Gouet, 2014) analysis reveals that the most similar crystal structure to *Tv*PPi-PFK is that of *Candidatus Prometheoarchaeum syntrophicum* PPi-PFK (*CPs*PPi-PFK, pdb entry 9CIR). The next closest is that of the Lyme disease spirochete *Borrelia burgdorferi* PPi-PFK (*Bb*PPi-PFK, pdb entry 1KZH) (Moore *et al*., 2002). These proteins share ∼29% sequence identity with *Tv*PPi-PFK. The carboxyl terminal domain of *Tv*PPi-PFK contains the modestly conserved GXXXR motif: GYELR, residues 307-311 in *Tv*PPi-PFK, GYEGR in 1KZH, and GHTMR in *Bb*PPi-PFK (Figure 4, Figure S.4). The catalytic conserved substrate-binding “MGR” motif of *Tv*PPi-PFK consists of amino acid residues 186-188. The substrate-binding site also includes the conserved catalytic Glu247 (Figure 4). ENDScript alignment reveals that the N-terminus domains of the proteins are better conserved than their C-terminus domains. The least conserved regions identified by ENDScript alignment are in the C-terminus domains of the three proteins (Figure 4). Compared to the C-terminus domains, the N-terminus domains are well-conserved except for an additional 14 amino acid insertion that extends the alpha helix 5 of *CPs*PPi-PFK (Figure 4). The ENDScript sausage plots and superposed protomers indicate that, despite the low (∼29%) sequence similarity, *Tv*PPi-PFK, *CPs*PPi-PFK, and *Bb*PPi-PFK have similar overall topologies (Figures 5a, 5b & 5c).

**Figure 4.**
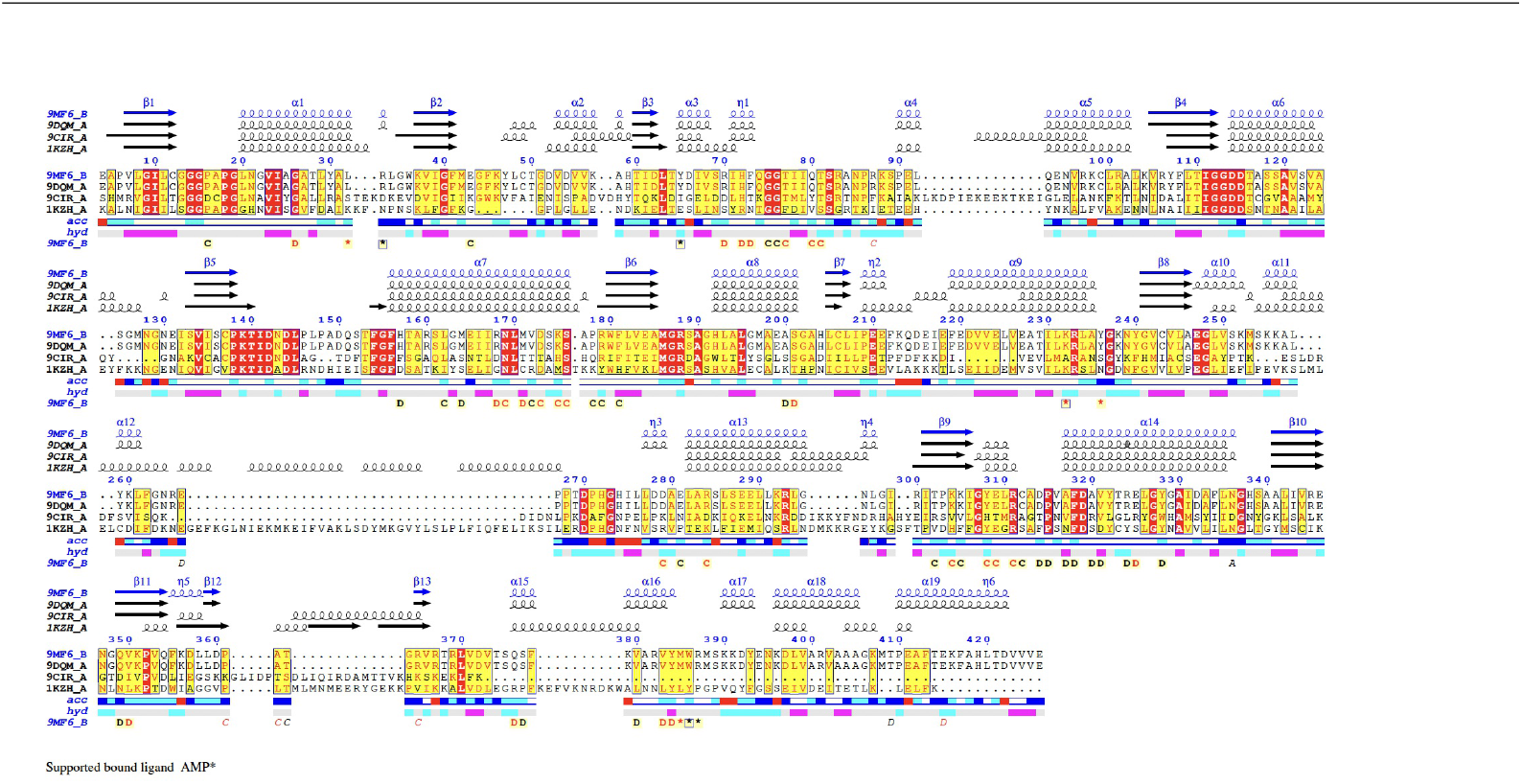
The amino-terminus domain of *Tv*PPi-PFK aligns well with those of its nearest structural neighbors. ENDScript structure-based-sequence alignment (Gouet *et al*., 2003, Gouet *et al*., 1999) reveals the nearest structural neighbors of *Tv*PPi-PFK as Lyme disease spirochete *Borrelia burgdorferi* PPi-PFK (*Bb*PPi-PFK, pdb entry 1KZH) (Moore *et al*., 2002) and *Candidatus Prometheoarchaeum syntrophicum* PPi-PFK (*CPs*PPi-PFK, pdb entry 9CIR). Also shown is another *Tv*PPi-PFK structure (pdb entry 9DQM). Identical and conserved residues are highlighted in red and yellow, respectively. The different secondary structure elements shown are alpha helices (*α*), 310-helices (*η*), beta strands (*β*), and beta turns (TT)(Gouet *et al*., 2003, Gouet *et al*., 1999)

**Figure 5.**
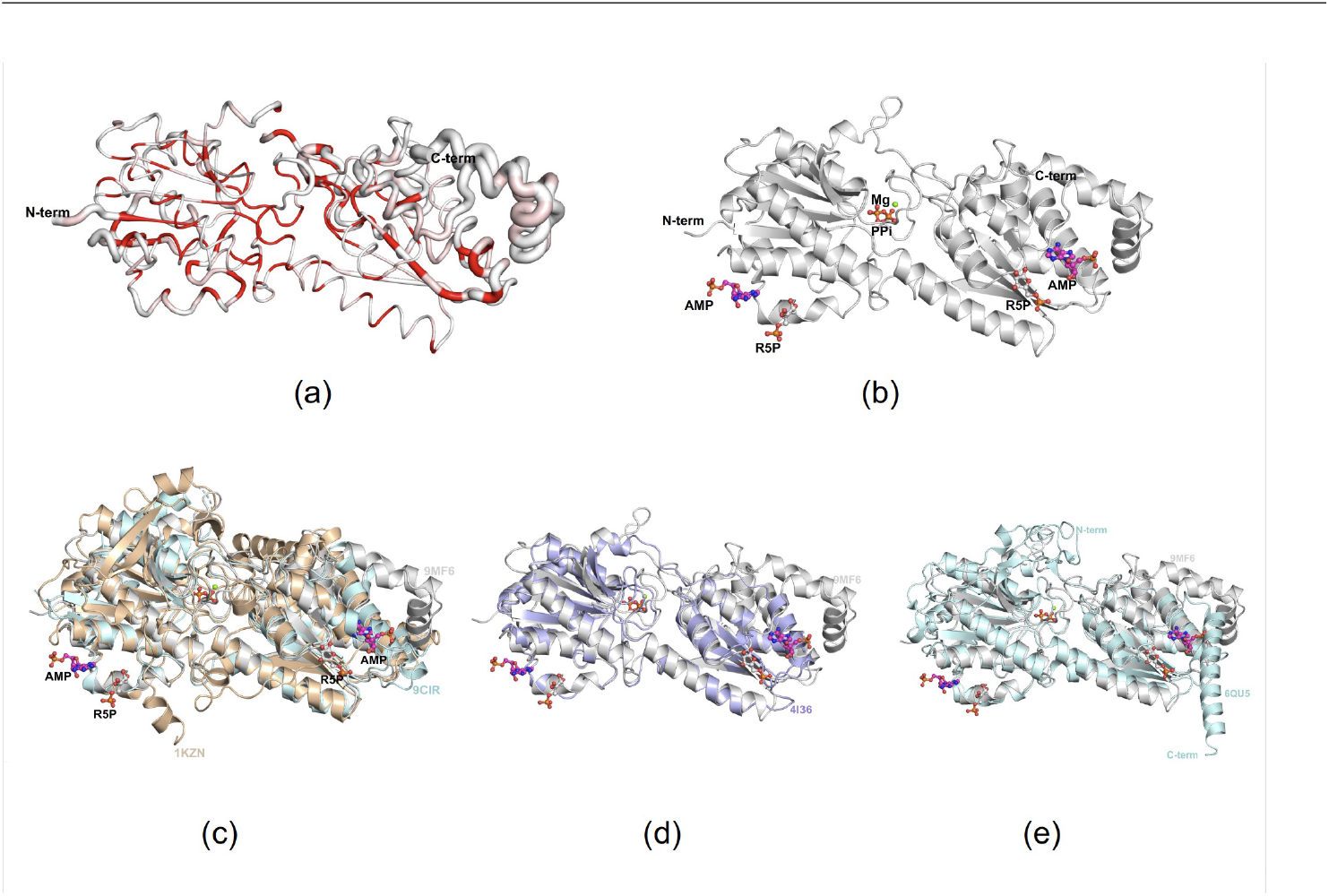
Comparison of *Tv*PPi-PFK to its structural neighbors. (a) Sausage plot calculated by ENDScript. The circumference of the ribbon (sausage) represents relative structural conservation of *Tv*PPi-PFK compared to *CPs*PPi-PFK and *Bb*PPi-PFK. Thinner ribbons represent regions with lower RMSD for alignment. In comparison, thicker ribbons represent higher RMSD for alignment, and the ribbons are colored by sequence conservation, with red indicating identical residues. (b) Same view of representative *Tv*PPi-PFK protomer (pdb entry 9MF6) showing Mg and PPi in the central pyrophosphate binding cavity at the interface of the two domains. Also shown are ligands at the AMP R5P at the dimer interface. (c) Superposed protomers of *Tv*PPi-PFK (pdb entry 9MF6, gray) superposed with *CPs*PPi-PFK (9CIR, green) and *Bb*PPi-PFK (1KZN, wheat) reveal the obstructed ATP-binding region in both *CPs*PPi-PFK and *Bb*PPi-PFK. (d) Superposed protomers of *Tv*PPi-PFK (pdb entry 9MF6, gray) and *Gst*ATP-PFK (4I36, purple) reveal conserved overall topology. (e) Likewise, superposed protomers of *Tv*PPi-PFK (pdb entry 9MF6, gray) and *Tb*ATP-PFK (6QU5, cyan) reveal overall structural similarity. Figures 5a-e are shown in the same orientation.

3D protein structure comparison against the entire RCSB protein database using the DALI server (http://ekhidna2.biocenter.helsinki.fi/dali) was also used to identify the closest structures to *Tv*PPi-PFK (Holm, 2022). DALI analyses agreed with ENDScript and identified *CPs*PPi-PFK (PDB entries 9CIT, 9CIS) as the closest structures to *Tv*PPi-PFK (Table S.1). Surprisingly, the next closest structure to *Tv*PPi-PFK that was identified was not another PPi-PFK but instead *Trypanosoma brucei* ATP-PFK (*Tb*ATP-PFK) (McNae *et al*., 2021). *Tb*ATP-PFK is topologically similar to *Tv*PPi-PFK despite having a larger size and amino and carboxyl termini extension (Figure 5d). Interestingly, allosteric inhibitors of *Tb*ATP-PFK have been identified that also clear *T. brucei* infections in mice (McNae *et al*., 2021). We plan future studies to test if these ligands can bind to *Tv*PPi-PFK or inhibit its activity. The *Tv*PPi-PFK protomer also shares topological similarity with the prototypical bacterial ATP-PFK from *Geobacillus stearothermophilus* (*Gst*ATP-PFK) (Mosser *et al*., 2013) (Figure 5e). The supplementary DALI alignment tables report RMSD values.

## Conclusion

We report three structure of *Tv*PPi-PFK deposited with the Worldwide PDB (wwPDB) under the accession code 9MDT (pdb_00009mdt), 9MF6 (pdb_00009mf6) and 9MED (pdb_00009med). *Tv*PPi-PFK is a cytosolic enzyme responsible for most (98%) of *T. vaginalis’* glycolytic flux and energetic needs (Rada *et al*., 2015) and the *Tv*PPi-PFK structures reveals inorganic pyrophosphate binding as needed for PPi-PFK enzymatic activity. Our structures reveal AMP and sugar phosphate binding at *Tv*PPi-PFK’s dimer interface. Mutagenesis and activity studies are planned to determine if these new AMP and sugar-phosphate binding sites are artefacts of crystallization or if they affect enzymatic activity.

## Supporting information

Supplementary Section

## Acknowledgements

We thank Dr. Lisa Tuttle for training and guidance with Mass Photometry experiments at The Hans Neurath Biophysics Core of the University of Washington Department of Biochemistry. We thank Abhishek Kancherla for technical assistance. This project is part of ongoing efforts to use SSGCID structures in undergraduate student training. This project has been funded in whole or in part with Federal funds from the National Institute of Allergy and Infectious Diseases, National Institutes of Health, Department of Health and Human Services, under Contract No.: 75N93022C00036. This research used resources the NYX beamline 19-ID, supported by the New York Structural Biology Center, at the National Synchrotron Light Source II, a U.S. Department of Energy (DOE) Office of Science User Facility operated for the DOE Office of Science by Brookhaven National Laboratory under Contract No. DE-SC0012704. The NYX detector instrumentation was supported by grant S10OD030394 through the Office of the Director of the National Institutes of Health. We are also grateful for start-up funds provided by the Dartmouth Cancer Center (OAA). The Dartmouth Cancer Center is supported by Cancer Center Support Grant; Award Number: P30CA023108 from the National Cancer Institute.

## Funding information

National Institute of Allergy and Infectious Diseases (contract No. 75N93022C00036 to Bart Staker); National Cancer Institute (contract No. P30CA023108 to Dragnev); U.S. Department of Energy, Office of Science (contract No. DE-SC0012704 to Brookhaven National Laboratory); National Institutes of Health, NIH Office of the Director (grant No. S10OD030394 to Brookhaven National Laboratory).

## Notes

### Competing Interest Statement

The authors have declared no competing interest.

### Summary of Updates

We have refined structures used in the figures and updated ligands. Author information updated

