## Supplementary Section for "Structural analyses of Trichomonas vaginalis pyrophosphate-dependent phosphofructokinase (TvPPi-PFK)"

Supplementary materials

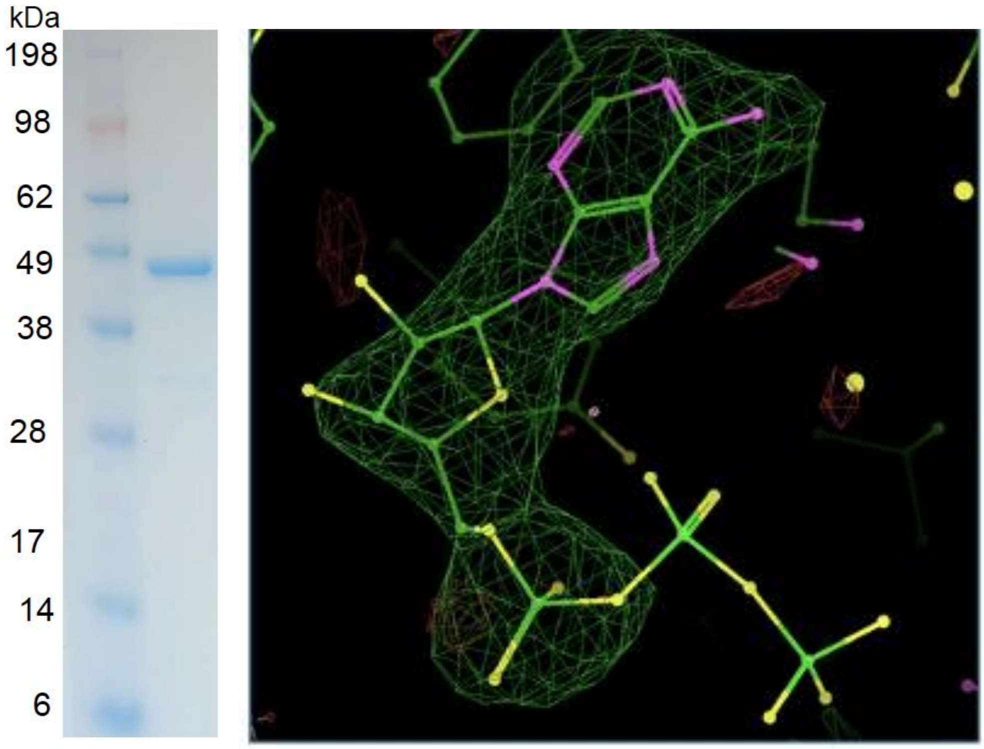


**(a)** **(b)**

**Figure S.1.** Protein purification and model fitting. (a) Simply Blue stained reduced NuPage gel of purified *Tv*PPi-PFK. (b) 3 sigma Fo-Fc omit reveal that ATP should not be modelled as there is no density for the PPi (9MDT). Instead AMP fits well into the maps. Similar quality maps were obtained for other ligands.

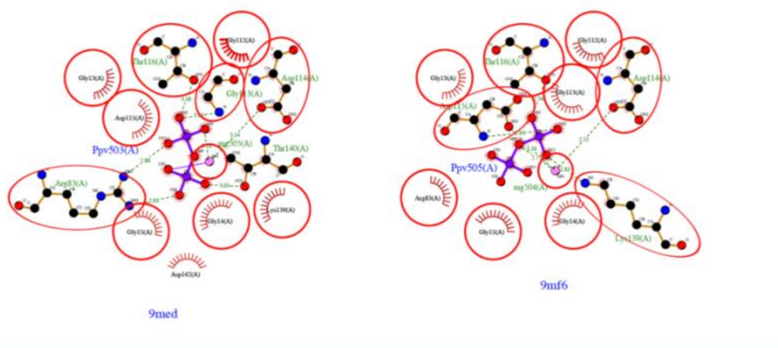


**(a)**

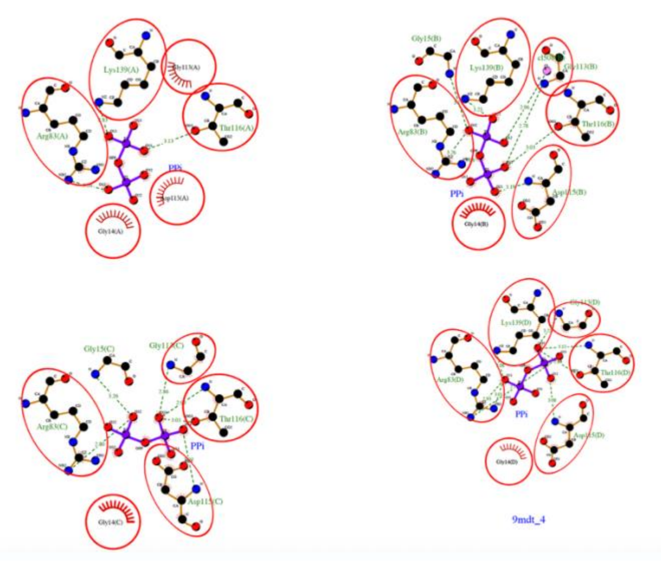


**(b)**

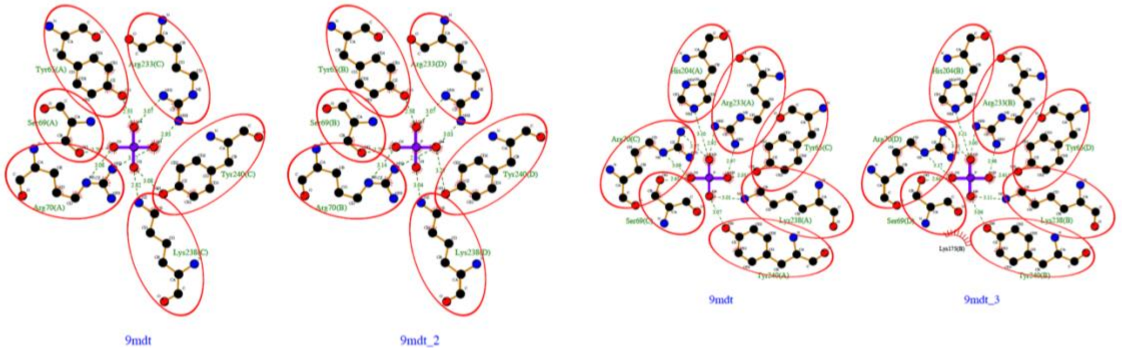


**(c)**

**Figure S.2.** PPi forms a network of interactions that includes key catalytic residues in the substrate binding cavity. (a) Comparison of ligand interaction plots generated with LIGPLOT+ showing amino acids involved in the *Tv*PPi-PFK's substrate-binding cavity of MgPPi$^{2-}$ complex (modelled as Mg and PPV). (b) PPi binding interaction plots reveal conserved amino acids involved for each *Tv*PPi-PFK protomer of pdb entry 9mdt. (c) Similarly, ligand interaction plots for inorganic phosphate (Pi) reveal conserved amino acids involved for each *Tv*PPi-PFK protomer of pdb entry 9mdt.

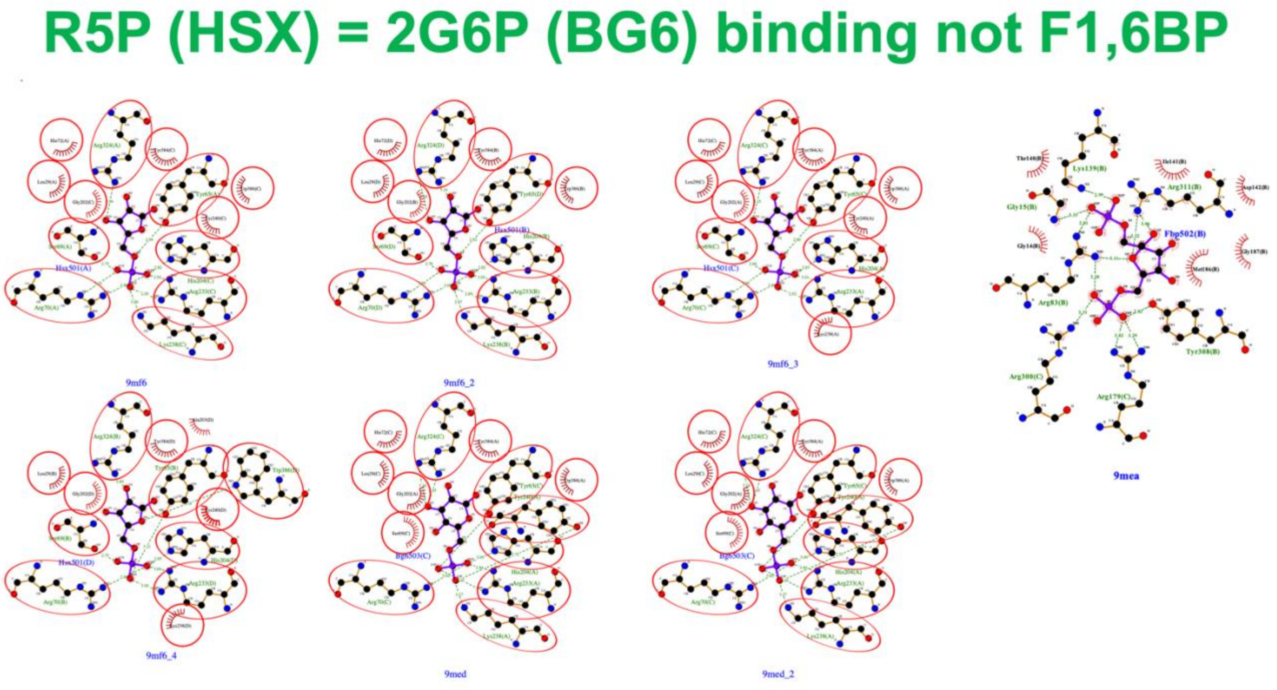


**Figure S.3.** Ribose-5-phosphate (R5P/HSX) and 2-glucose-6-phosphate (2G6P/BG6) bind in a unique sugar-phosphate binding cavity at the dimer interface that forms a network of shared amino acid interactions shown in ligand interaction plots generated with LIGPLOT+. The R5P/2G6P binding site is distinct from the *Tv*PPi-PFK catalytic site occupied by fructose-6-phosphate as exemplified by pdb entry 9MEA.

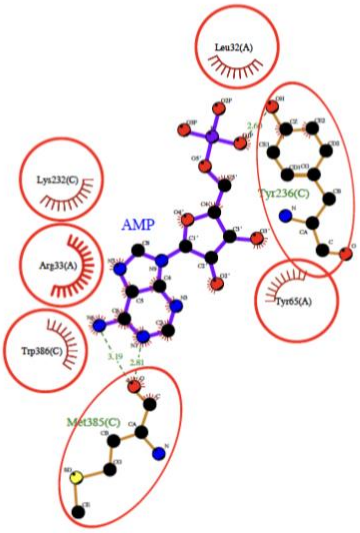


9mdt_4

**Figure S.4.** Ligand interaction plots generated with LIGPLOT+ reveal identical residues involved in AMP binding across the dimer interface of *Tv*PPi-PFK.

**Figure S.5.** Additional ENDScript structure-based-sequence alignment plots using different chains reveal the nearest structural neighbors of *Tv*PPi-PFK as Lyme disease spirochete *Borrelia burgdorferi* PPi-PFK (*Bb*PPi-PFK, pdb entry 1KZH)(Moore *et al.*, 2002) and *Candidatus Prometheoarchaeum syntrophicum* PPi-PFK (*CPs*PPi-PFK, pdb entry 9CIR). Also shown is another *Tv*PPi-PFK structure (pdb entry 9DQM). Identical and conserved residues are highlighted in red and yellow, respectively. The different secondary structure elements shown are alpha helices (α), 310-helices (η), beta strands (β), and beta turns (TT)(Gouet *et al.*, 2003, Gouet *et al.*, 1999).

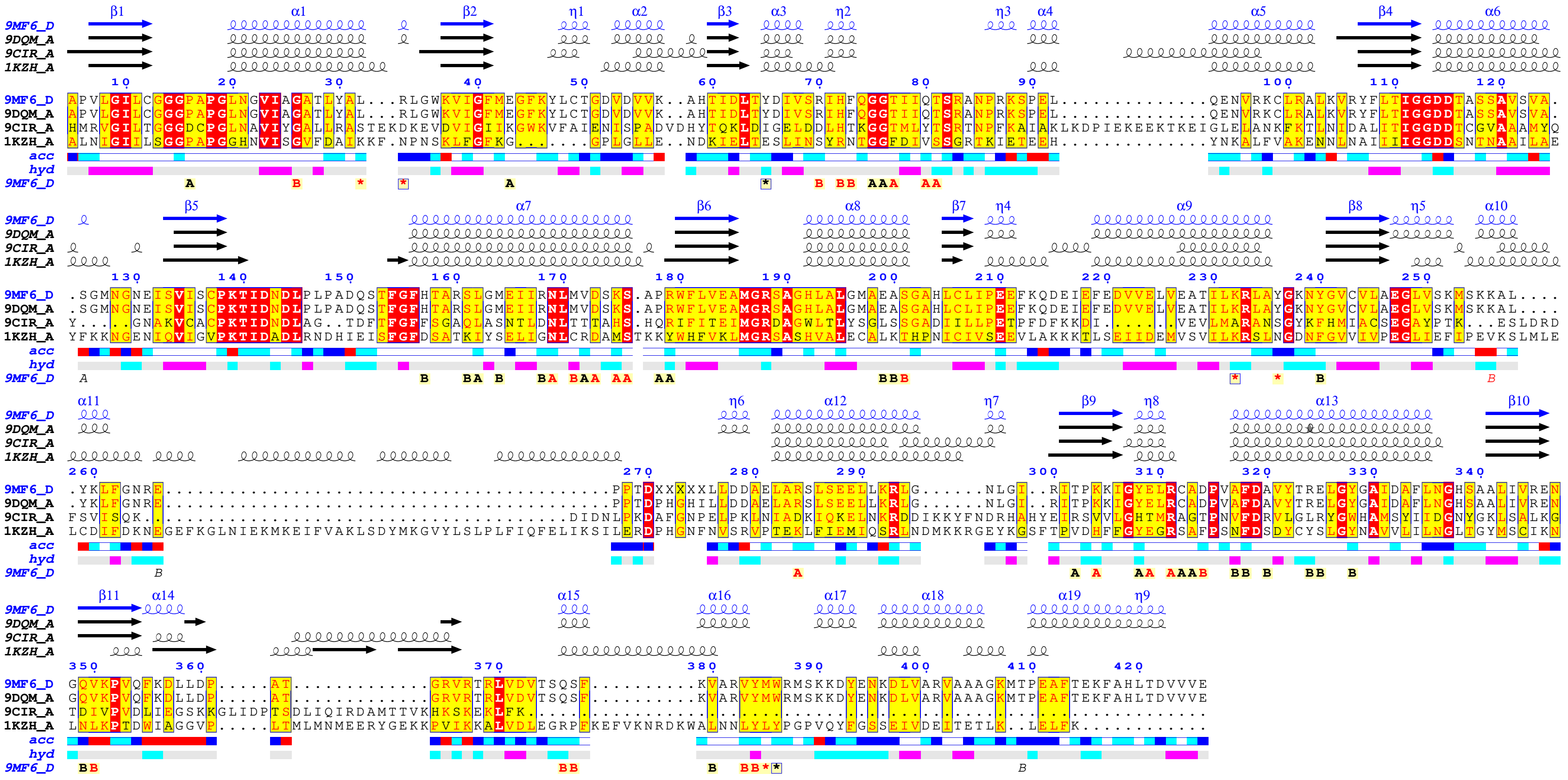


Supported bound ligand AMP*

Figure S.4a

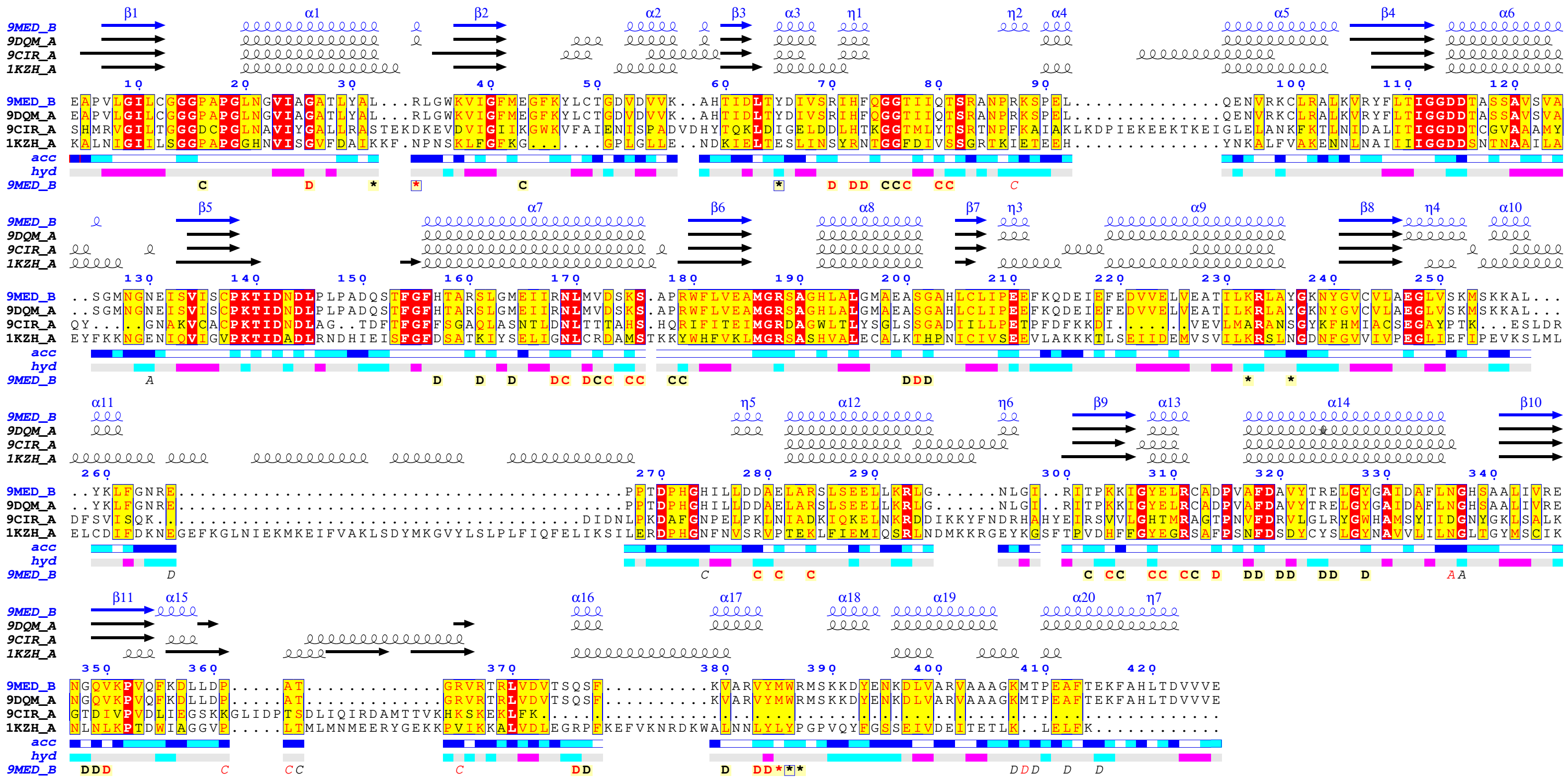


Supported bound ligand AMP*

Figure S.4b

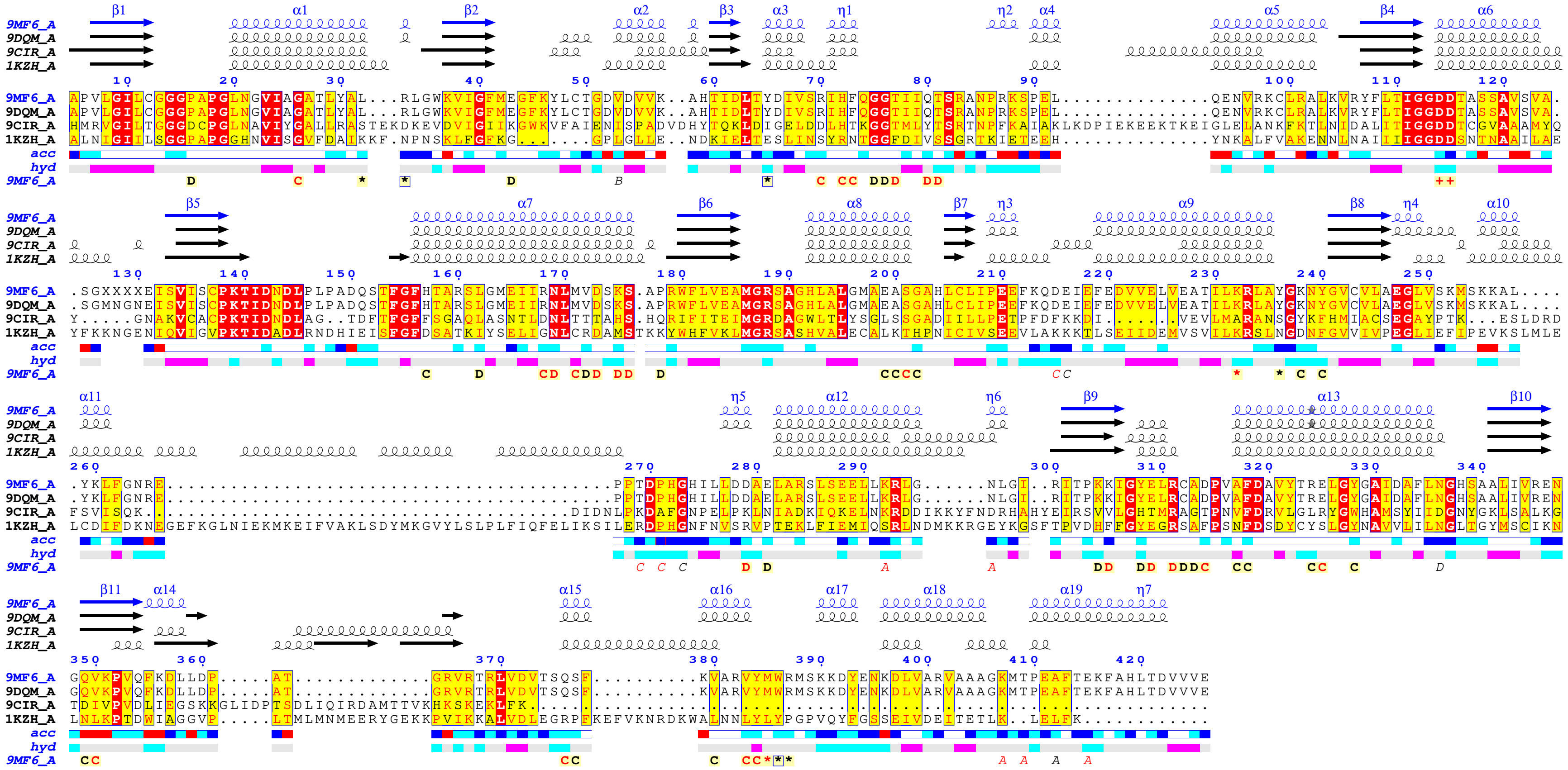


Supported bound ligands AMP* MG+

Figure S.4c

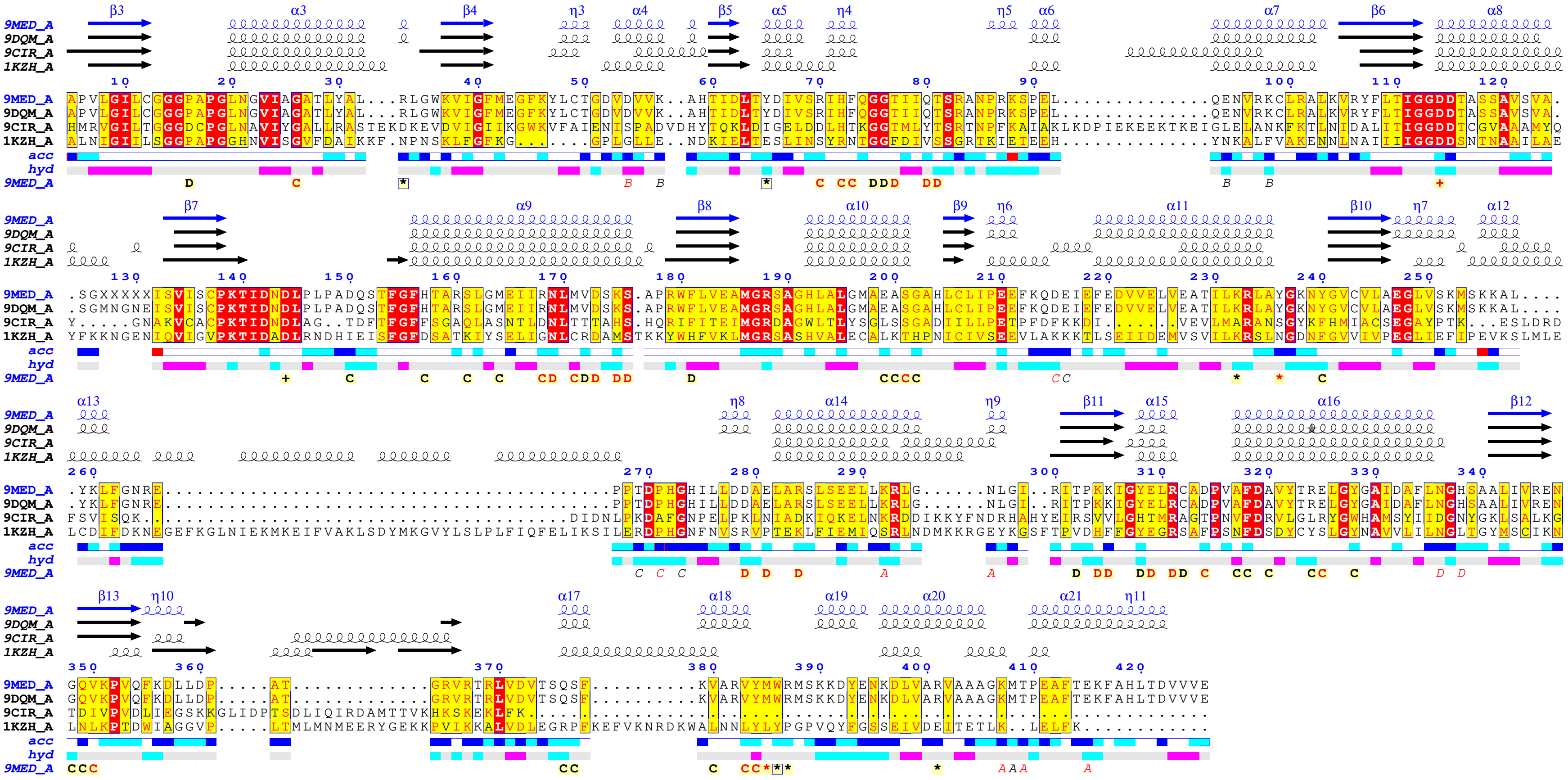

Supported bound ligands AMP* MG+

Figure S.4d

### Results: 9mf6A

#### Query: 9mf6A

MOLECULE: PYROPHOSPHATE--FRUCTOSE 6-PHOSPHATE 1-PHOSPHOTRAN

Select neighbours (check boxes) for viewing as multiple structural alignment or 3D superimposition. The list of neighbours is sorted by Z-score. Similarities with a Z-score lower than 2 are spurious. Each neighbour has links to pairwise structural alignment with the query structure, and to the PDB format coordinate file where the neighbour is superimposed onto the query structure.

Structural Alignment ☑ Expand gaps 3D Superimposition (PV) SANS PANZ Pfam Reset Selection

#### Summary

| No: | Chain | Z | rmsd | lali | nres | %id | PDB | Description |
|---|---|---|---|---|---|---|---|---|
| 1: | 9mf6-A | 68.6 | 0.0 | 418 | 418 | 100 | PDB | MOLECULE: PYROPHOSPHATE--FRUCTOSE 6-PHOSPHATE 1-PHOSPHOTRAN |
| 2: | 9med-A | 64.8 | 0.3 | 417 | 417 | 100 | PDB | MOLECULE: PYROPHOSPHATE--FRUCTOSE 6-PHOSPHATE 1-PHOSPHOTRAN |
| 3: | 9mdt-A | 64.7 | 0.3 | 417 | 417 | 100 | PDB | MOLECULE: PYROPHOSPHATE--FRUCTOSE 6-PHOSPHATE 1-PHOSPHOTRAN |
| 4: | 9mea-A | 64.5 | 0.4 | 418 | 418 | 100 | PDB | MOLECULE: PYROPHOSPHATE--FRUCTOSE 6-PHOSPHATE 1-PHOSPHOTRAN |
| 5: | 9mec-A | 64.4 | 0.3 | 418 | 422 | 100 | PDB | MOLECULE: PYROPHOSPHATE--FRUCTOSE 6-PHOSPHATE 1-PHOSPHOTRAN |
| 6: | 9mea-B | 60.8 | 0.9 | 418 | 423 | 100 | PDB | MOLECULE: PYROPHOSPHATE--FRUCTOSE 6-PHOSPHATE 1-PHOSPHOTRAN |
| 7: | 9dqm-B | 60.5 | 1.1 | 418 | 421 | 100 | PDB | MOLECULE: PYROPHOSPHATE--FRUCTOSE 6-PHOSPHATE 1-PHOSPHOTRAN |
| 8: | 9mec-B | 60.3 | 1.0 | 418 | 423 | 100 | PDB | MOLECULE: PYROPHOSPHATE--FRUCTOSE 6-PHOSPHATE 1-PHOSPHOTRAN |
| 9: | 9med-B | 60.3 | 1.0 | 418 | 423 | 100 | PDB | MOLECULE: PYROPHOSPHATE--FRUCTOSE 6-PHOSPHATE 1-PHOSPHOTRAN |
| 10: | 9mf6-B | 60.2 | 1.0 | 418 | 423 | 100 | PDB | MOLECULE: PYROPHOSPHATE--FRUCTOSE 6-PHOSPHATE 1-PHOSPHOTRAN |
| 11: | 9mdt-B | 60.2 | 1.0 | 418 | 423 | 100 | PDB | MOLECULE: PYROPHOSPHATE--FRUCTOSE 6-PHOSPHATE 1-PHOSPHOTRAN |
| 12: | 9dqm-A | 59.9 | 1.3 | 418 | 420 | 100 | PDB | MOLECULE: PYROPHOSPHATE--FRUCTOSE 6-PHOSPHATE 1-PHOSPHOTRAN |
| 13: | 9mea-D | 59.9 | 1.1 | 413 | 417 | 100 | PDB | MOLECULE: PYROPHOSPHATE--FRUCTOSE 6-PHOSPHATE 1-PHOSPHOTRAN |
| 14: | 9mdt-D | 59.8 | 1.2 | 413 | 417 | 100 | PDB | MOLECULE: PYROPHOSPHATE--FRUCTOSE 6-PHOSPHATE 1-PHOSPHOTRAN |
| 15: | 9mec-D | 59.8 | 1.1 | 413 | 417 | 100 | PDB | MOLECULE: PYROPHOSPHATE--FRUCTOSE 6-PHOSPHATE 1-PHOSPHOTRAN |
| 16: | 9mdt-C | 59.4 | 1.3 | 418 | 423 | 100 | PDB | MOLECULE: PYROPHOSPHATE--FRUCTOSE 6-PHOSPHATE 1-PHOSPHOTRAN |
| 17: | 9mf6-D | 59.3 | 1.2 | 413 | 417 | 100 | PDB | MOLECULE: PYROPHOSPHATE--FRUCTOSE 6-PHOSPHATE 1-PHOSPHOTRAN |
| 18: | 9mea-C | 59.3 | 1.3 | 418 | 423 | 100 | PDB | MOLECULE: PYROPHOSPHATE--FRUCTOSE 6-PHOSPHATE 1-PHOSPHOTRAN |
| 19: | 9med-D | 59.3 | 1.2 | 413 | 417 | 100 | PDB | MOLECULE: PYROPHOSPHATE--FRUCTOSE 6-PHOSPHATE 1-PHOSPHOTRAN |
| 20: | 9mec-C | 59.2 | 1.3 | 418 | 423 | 100 | PDB | MOLECULE: PYROPHOSPHATE--FRUCTOSE 6-PHOSPHATE 1-PHOSPHOTRAN |
| 21: | 9med-C | 59.1 | 1.3 | 418 | 423 | 100 | PDB | MOLECULE: PYROPHOSPHATE--FRUCTOSE 6-PHOSPHATE 1-PHOSPHOTRAN |
| 22: | 9mf6-C | 58.8 | 1.4 | 415 | 420 | 100 | PDB | MOLECULE: PYROPHOSPHATE--FRUCTOSE 6-PHOSPHATE 1-PHOSPHOTRAN |
| 23: | 9cit-D | 39.9 | 2.7 | 357 | 378 | 31 | PDB | MOLECULE: 6-PHOSPHOFRUCTOKINASE; |
| 24: | 9cir-D | 39.4 | 2.6 | 350 | 372 | 31 | PDB | MOLECULE: 6-PHOSPHOFRUCTOKINASE; |
| 25: | 9cis-C | 39.3 | 2.6 | 355 | 377 | 30 | PDB | MOLECULE: 6-PHOSPHOFRUCTOKINASE; |
| 26: | 9cir-C | 39.2 | 2.7 | 352 | 377 | 30 | PDB | MOLECULE: 6-PHOSPHOFRUCTOKINASE; |
| 27: | 9cit-C | 39.1 | 2.7 | 353 | 380 | 30 | PDB | MOLECULE: 6-PHOSPHOFRUCTOKINASE; |
| 28: | 9cis-A | 39.0 | 2.9 | 360 | 384 | 31 | PDB | MOLECULE: 6-PHOSPHOFRUCTOKINASE; |
| 29: | 9cir-B | 38.9 | 3.1 | 363 | 391 | 30 | PDB | MOLECULE: 6-PHOSPHOFRUCTOKINASE; |
| 30: | 9cit-B | 38.8 | 2.8 | 357 | 385 | 31 | PDB | MOLECULE: 6-PHOSPHOFRUCTOKINASE; |
| 31: | 9cir-A | 38.8 | 2.8 | 356 | 385 | 30 | PDB | MOLECULE: 6-PHOSPHOFRUCTOKINASE; |
| 32: | 9cit-A | 38.6 | 2.9 | 360 | 389 | 30 | PDB | MOLECULE: 6-PHOSPHOFRUCTOKINASE; |
| 33: | 9cis-B | 38.4 | 2.7 | 350 | 375 | 31 | PDB | MOLECULE: 6-PHOSPHOFRUCTOKINASE; |
| 34: | 9cis-D | 38.4 | 2.6 | 347 | 366 | 31 | PDB | MOLECULE: 6-PHOSPHOFRUCTOKINASE; |
| 35: | 6qu4-B | 37.9 | 2.5 | 337 | 439 | 29 | PDB | MOLECULE: ATP-DEPENDENT 6-PHOSPHOFRUCTOKINASE; |
| 36: | 6qu5-F | 37.8 | 2.5 | 341 | 456 | 29 | PDB | MOLECULE: ATP-DEPENDENT 6-PHOSPHOFRUCTOKINASE; |
| 37: | 3pfk-A | 37.3 | 2.2 | 318 | 319 | 27 | PDB | MOLECULE: PHOSPHOFRUCTOKINASE; |
| 38: | 6qu3-D | 37.2 | 2.5 | 347 | 461 | 29 | PDB | MOLECULE: ATP-DEPENDENT 6-PHOSPHOFRUCTOKINASE; |
| 39: | 6qu3-B | 37.1 | 2.3 | 339 | 446 | 29 | PDB | MOLECULE: ATP-DEPENDENT 6-PHOSPHOFRUCTOKINASE; |
| 40: | 1mto-G | 37.0 | 2.0 | 317 | 319 | 27 | PDB | MOLECULE: 6-PHOSPHOFRUCTOKINASE; |
| 41: | 1pfk-A | 37.0 | 2.1 | 317 | 320 | 27 | PDB | MOLECULE: PHOSPHOFRUCTOKINASE; |
| 42: | 1mto-A | 37.0 | 2.0 | 316 | 319 | 27 | PDB | MOLECULE: 6-PHOSPHOFRUCTOKINASE; |
| 43: | 4pfk-A | 36.9 | 2.2 | 318 | 319 | 27 | PDB | MOLECULE: PHOSPHOFRUCTOKINASE; |
| 44: | 3u39-A | 36.9 | 2.1 | 317 | 319 | 27 | PDB | MOLECULE: 6-PHOSPHOFRUCTOKINASE; |
| 45: | 3u39-D | 36.9 | 2.1 | 317 | 319 | 27 | PDB | MOLECULE: 6-PHOSPHOFRUCTOKINASE; |
| 46: | 2hig-B | 36.9 | 2.0 | 332 | 422 | 30 | PDB | MOLECULE: 6-PHOSPHO-1-FRUCTOKINASE; |
| 47: | 5xz7-A | 36.8 | 2.2 | 317 | 322 | 26 | PDB | MOLECULE: ATP-DEPENDENT 6-PHOSPHOFRUCTOKINASE; |
| 48: | 1mto-D | 36.8 | 2.1 | 318 | 319 | 27 | PDB | MOLECULE: 6-PHOSPHOFRUCTOKINASE; |
| 49: | 4i4i-D | 36.7 | 2.1 | 316 | 319 | 27 | PDB | MOLECULE: 6-PHOSPHOFRUCTOKINASE; |
| 50: | 1mto-C | 36.7 | 2.0 | 317 | 319 | 27 | PDB | MOLECULE: 6-PHOSPHOFRUCTOKINASE; |
| 51: | 3u39-C | 36.7 | 2.2 | 318 | 319 | 27 | PDB | MOLECULE: 6-PHOSPHOFRUCTOKINASE; |
| 52: | 4a3s-B | 36.7 | 2.2 | 317 | 319 | 25 | PDB | MOLECULE: 6-PHOSPHOFRUCTOKINASE; |
| 53: | 5xz8-A | 36.7 | 2.0 | 313 | 317 | 26 | PDB | MOLECULE: ATP-DEPENDENT 6-PHOSPHOFRUCTOKINASE; |
| 54: | 1mto-B | 36.6 | 2.1 | 318 | 319 | 27 | PDB | MOLECULE: 6-PHOSPHOFRUCTOKINASE; |
| 55: | 6qu5-A | 36.6 | 2.4 | 338 | 452 | 29 | PDB | MOLECULE: ATP-DEPENDENT 6-PHOSPHOFRUCTOKINASE; |
| 56: | 3u39-B | 36.6 | 2.1 | 317 | 318 | 27 | PDB | MOLECULE: 6-PHOSPHOFRUCTOKINASE; |
| 57: | 6qu3-C | 36.6 | 2.7 | 340 | 446 | 29 | PDB | MOLECULE: ATP-DEPENDENT 6-PHOSPHOFRUCTOKINASE; |
| 58: | 1mto-F | 36.6 | 2.1 | 318 | 319 | 27 | PDB | MOLECULE: 6-PHOSPHOFRUCTOKINASE; |
| 59: | 4i36-D | 36.6 | 2.1 | 316 | 319 | 27 | PDB | MOLECULE: 6-PHOSPHOFRUCTOKINASE; |
| 60: | 4a3s-A | 36.6 | 2.2 | 317 | 319 | 25 | PDB | MOLECULE: 6-PHOSPHOFRUCTOKINASE; |

### Results: 9mdtC

#### Query: 9mdtC

MOLECULE: PYROPHOSPHATE--FRUCTOSE 6-PHOSPHATE 1-PHOSPHOTRAN

Select neighbours (check boxes) for viewing as multiple structural alignment or 3D superimposition. The list of neighbours is sorted by Z-score. Similarities with a Z-score lower than 2 are spurious. Each neighbour has links to pairwise structural alignment with the query structure, and to the PDB format coordinate file where the neighbour is superimposed onto the query structure.

Structural Alignment ☑ Expand gaps 3D Superimposition (PV) SANS PANZ Pfam Reset Selection

#### Summary

| No: | Chain | Z | rmsd | lali | nres | %id | PDB | Description |
|---|---|---|---|---|---|---|---|---|
| 1: | 9mdt-C | 66.7 | 0.0 | 423 | 423 | 100 | PDB | MOLECULE: PYROPHOSPHATE--FRUCTOSE 6-PHOSPHATE 1-PHOSPHOTRAN |
| 2: | 9mec-C | 64.9 | 0.1 | 423 | 423 | 100 | PDB | MOLECULE: PYROPHOSPHATE--FRUCTOSE 6-PHOSPHATE 1-PHOSPHOTRAN |
| 3: | 9mea-C | 64.7 | 0.1 | 423 | 423 | 100 | PDB | MOLECULE: PYROPHOSPHATE--FRUCTOSE 6-PHOSPHATE 1-PHOSPHOTRAN |
| 4: | 9med-C | 64.6 | 0.2 | 423 | 423 | 100 | PDB | MOLECULE: PYROPHOSPHATE--FRUCTOSE 6-PHOSPHATE 1-PHOSPHOTRAN |
| 5: | 9mdt-D | 64.0 | 0.3 | 417 | 417 | 100 | PDB | MOLECULE: PYROPHOSPHATE--FRUCTOSE 6-PHOSPHATE 1-PHOSPHOTRAN |
| 6: | 9mec-D | 63.8 | 0.3 | 417 | 417 | 100 | PDB | MOLECULE: PYROPHOSPHATE--FRUCTOSE 6-PHOSPHATE 1-PHOSPHOTRAN |
| 7: | 9mf6-C | 63.5 | 0.3 | 420 | 420 | 100 | PDB | MOLECULE: PYROPHOSPHATE--FRUCTOSE 6-PHOSPHATE 1-PHOSPHOTRAN |
| 8: | 9mdt-B | 63.5 | 0.5 | 423 | 423 | 100 | PDB | MOLECULE: PYROPHOSPHATE--FRUCTOSE 6-PHOSPHATE 1-PHOSPHOTRAN |
| 9: | 9med-D | 63.4 | 0.3 | 417 | 417 | 100 | PDB | MOLECULE: PYROPHOSPHATE--FRUCTOSE 6-PHOSPHATE 1-PHOSPHOTRAN |
| 10: | 9mec-B | 63.3 | 0.5 | 423 | 423 | 100 | PDB | MOLECULE: PYROPHOSPHATE--FRUCTOSE 6-PHOSPHATE 1-PHOSPHOTRAN |
| 11: | 9mea-D | 63.1 | 0.4 | 417 | 417 | 100 | PDB | MOLECULE: PYROPHOSPHATE--FRUCTOSE 6-PHOSPHATE 1-PHOSPHOTRAN |
| 12: | 9med-B | 62.9 | 0.5 | 423 | 423 | 100 | PDB | MOLECULE: PYROPHOSPHATE--FRUCTOSE 6-PHOSPHATE 1-PHOSPHOTRAN |
| 13: | 9mf6-D | 62.5 | 0.4 | 417 | 417 | 100 | PDB | MOLECULE: PYROPHOSPHATE--FRUCTOSE 6-PHOSPHATE 1-PHOSPHOTRAN |
| 14: | 9mf6-B | 62.5 | 0.6 | 423 | 423 | 100 | PDB | MOLECULE: PYROPHOSPHATE--FRUCTOSE 6-PHOSPHATE 1-PHOSPHOTRAN |
| 15: | 9mea-B | 62.3 | 0.6 | 423 | 423 | 100 | PDB | MOLECULE: PYROPHOSPHATE--FRUCTOSE 6-PHOSPHATE 1-PHOSPHOTRAN |
| 16: | 9dqm-B | 62.3 | 0.6 | 421 | 421 | 100 | PDB | MOLECULE: PYROPHOSPHATE--FRUCTOSE 6-PHOSPHATE 1-PHOSPHOTRAN |
| 17: | 9dqm-A | 62.1 | 0.6 | 420 | 420 | 100 | PDB | MOLECULE: PYROPHOSPHATE--FRUCTOSE 6-PHOSPHATE 1-PHOSPHOTRAN |
| 18: | 9mdt-A | 60.6 | 1.1 | 417 | 417 | 100 | PDB | MOLECULE: PYROPHOSPHATE--FRUCTOSE 6-PHOSPHATE 1-PHOSPHOTRAN |
| 19: | 9mec-A | 60.5 | 1.2 | 422 | 422 | 100 | PDB | MOLECULE: PYROPHOSPHATE--FRUCTOSE 6-PHOSPHATE 1-PHOSPHOTRAN |
| 20: | 9med-A | 60.3 | 1.2 | 417 | 417 | 100 | PDB | MOLECULE: PYROPHOSPHATE--FRUCTOSE 6-PHOSPHATE 1-PHOSPHOTRAN |
| 21: | 9mea-A | 60.2 | 1.2 | 418 | 418 | 100 | PDB | MOLECULE: PYROPHOSPHATE--FRUCTOSE 6-PHOSPHATE 1-PHOSPHOTRAN |
| 22: | 9mf6-A | 59.4 | 1.3 | 418 | 418 | 100 | PDB | MOLECULE: PYROPHOSPHATE--FRUCTOSE 6-PHOSPHATE 1-PHOSPHOTRAN |
| 23: | 9cit-D | 37.4 | 2.7 | 357 | 378 | 31 | PDB | MOLECULE: 6-PHOSPHOFRUCTOKINASE; |
| 24: | 9cir-C | 37.0 | 2.8 | 353 | 377 | 31 | PDB | MOLECULE: 6-PHOSPHOFRUCTOKINASE; |
| 25: | 9cis-C | 37.0 | 2.7 | 356 | 377 | 31 | PDB | MOLECULE: 6-PHOSPHOFRUCTOKINASE; |
| 26: | 9cir-D | 37.0 | 2.8 | 352 | 372 | 31 | PDB | MOLECULE: 6-PHOSPHOFRUCTOKINASE; |
| 27: | 9cir-A | 36.5 | 2.8 | 356 | 385 | 30 | PDB | MOLECULE: 6-PHOSPHOFRUCTOKINASE; |
| 28: | 9cit-C | 36.5 | 2.8 | 352 | 380 | 31 | PDB | MOLECULE: 6-PHOSPHOFRUCTOKINASE; |
| 29: | 9cis-A | 36.4 | 2.9 | 360 | 384 | 31 | PDB | MOLECULE: 6-PHOSPHOFRUCTOKINASE; |
| 30: | 9cis-B | 36.4 | 2.8 | 350 | 375 | 31 | PDB | MOLECULE: 6-PHOSPHOFRUCTOKINASE; |
| 31: | 9cit-B | 36.3 | 2.8 | 355 | 385 | 31 | PDB | MOLECULE: 6-PHOSPHOFRUCTOKINASE; |
| 32: | 9cir-B | 36.3 | 3.0 | 362 | 391 | 31 | PDB | MOLECULE: 6-PHOSPHOFRUCTOKINASE; |
| 33: | 6qu5-F | 36.3 | 2.6 | 346 | 456 | 29 | PDB | MOLECULE: ATP-DEPENDENT 6-PHOSPHOFRUCTOKINASE; |
| 34: | 9cis-D | 35.9 | 2.7 | 345 | 366 | 31 | PDB | MOLECULE: 6-PHOSPHOFRUCTOKINASE; |
| 35: | 9cit-A | 35.8 | 2.9 | 358 | 389 | 30 | PDB | MOLECULE: 6-PHOSPHOFRUCTOKINASE; |
| 36: | 5xz7-A | 35.6 | 2.0 | 318 | 322 | 25 | PDB | MOLECULE: ATP-DEPENDENT 6-PHOSPHOFRUCTOKINASE; |
| 37: | 6qu3-D | 35.5 | 2.5 | 351 | 461 | 28 | PDB | MOLECULE: ATP-DEPENDENT 6-PHOSPHOFRUCTOKINASE; |
| 38: | 3pfk-A | 35.4 | 2.0 | 317 | 319 | 27 | PDB | MOLECULE: PHOSPHOFRUCTOKINASE; |
| 39: | 1pfk-B | 35.3 | 1.9 | 315 | 320 | 27 | PDB | MOLECULE: PHOSPHOFRUCTOKINASE; |
| 40: | 5xz8-A | 35.3 | 1.9 | 313 | 317 | 26 | PDB | MOLECULE: ATP-DEPENDENT 6-PHOSPHOFRUCTOKINASE; |
| 41: | 4a3s-A | 35.2 | 1.9 | 315 | 319 | 25 | PDB | MOLECULE: 6-PHOSPHOFRUCTOKINASE; |
| 42: | 4a3s-B | 35.2 | 1.9 | 315 | 319 | 25 | PDB | MOLECULE: 6-PHOSPHOFRUCTOKINASE; |
| 43: | 8w2g-A | 35.0 | 3.3 | 341 | 741 | 24 | PDB | MOLECULE: ATP-DEPENDENT 6-PHOSPHOFRUCTOKINASE, LIVER TYPE; |
| 44: | 1mto-G | 35.0 | 1.9 | 315 | 319 | 27 | PDB | MOLECULE: 6-PHOSPHOFRUCTOKINASE; |
| 45: | 1pfk-A | 35.0 | 2.1 | 317 | 320 | 27 | PDB | MOLECULE: PHOSPHOFRUCTOKINASE; |
| 46: | 8w2g-B | 35.0 | 3.4 | 342 | 741 | 24 | PDB | MOLECULE: ATP-DEPENDENT 6-PHOSPHOFRUCTOKINASE, LIVER TYPE; |

### Results: 9medB

#### Query: 9medB

MOLECULE: PYROPHOSPHATE--FRUCTOSE 6-PHOSPHATE 1-PHOSPHOTRAN

Select neighbours (check boxes) for viewing as multiple structural alignment or 3D superimposition. The list of neighbours is sorted by Z-score. Similarities with a Z-score lower than 2 are spurious. Each neighbour has links to pairwise structural alignment with the query structure, and to the PDB format coordinate file where the neighbour is superimposed onto the query structure.

Structural Alignment ☑ Expand gaps 3D Superimposition (PV) SANS PANZ Pfam Reset Selection

#### Summary

| No: | Chain | Z | rmsd | lali | nres | %id | PDB | Description |
|---|---|---|---|---|---|---|---|---|
| 1: | 9med-B | 67.0 | 0.0 | 423 | 423 | 100 | PDB | MOLECULE: PYROPHOSPHATE--FRUCTOSE 6-PHOSPHATE 1-PHOSPHOTRAN |
| 2: | 9mec-B | 65.0 | 0.1 | 423 | 423 | 100 | PDB | MOLECULE: PYROPHOSPHATE--FRUCTOSE 6-PHOSPHATE 1-PHOSPHOTRAN |
| 3: | 9mdt-B | 64.5 | 0.2 | 423 | 423 | 100 | PDB | MOLECULE: PYROPHOSPHATE--FRUCTOSE 6-PHOSPHATE 1-PHOSPHOTRAN |
| 4: | 9mea-B | 64.2 | 0.3 | 423 | 423 | 100 | PDB | MOLECULE: PYROPHOSPHATE--FRUCTOSE 6-PHOSPHATE 1-PHOSPHOTRAN |
| 5: | 9mf6-B | 63.8 | 0.3 | 423 | 423 | 100 | PDB | MOLECULE: PYROPHOSPHATE--FRUCTOSE 6-PHOSPHATE 1-PHOSPHOTRAN |
| 6: | 9mdt-D | 63.4 | 0.4 | 417 | 417 | 100 | PDB | MOLECULE: PYROPHOSPHATE--FRUCTOSE 6-PHOSPHATE 1-PHOSPHOTRAN |
| 7: | 9mea-D | 63.4 | 0.3 | 417 | 417 | 100 | PDB | MOLECULE: PYROPHOSPHATE--FRUCTOSE 6-PHOSPHATE 1-PHOSPHOTRAN |
| 8: | 9mec-D | 63.3 | 0.4 | 417 | 417 | 100 | PDB | MOLECULE: PYROPHOSPHATE--FRUCTOSE 6-PHOSPHATE 1-PHOSPHOTRAN |
| 9: | 9med-D | 63.1 | 0.4 | 417 | 417 | 100 | PDB | MOLECULE: PYROPHOSPHATE--FRUCTOSE 6-PHOSPHATE 1-PHOSPHOTRAN |
| 10: | 9mec-C | 63.0 | 0.6 | 423 | 423 | 100 | PDB | MOLECULE: PYROPHOSPHATE--FRUCTOSE 6-PHOSPHATE 1-PHOSPHOTRAN |
| 11: | 9mea-C | 62.9 | 0.5 | 423 | 423 | 100 | PDB | MOLECULE: PYROPHOSPHATE--FRUCTOSE 6-PHOSPHATE 1-PHOSPHOTRAN |
| 12: | 9med-C | 62.9 | 0.6 | 423 | 423 | 100 | PDB | MOLECULE: PYROPHOSPHATE--FRUCTOSE 6-PHOSPHATE 1-PHOSPHOTRAN |
| 13: | 9mdt-C | 62.9 | 0.5 | 423 | 423 | 100 | PDB | MOLECULE: PYROPHOSPHATE--FRUCTOSE 6-PHOSPHATE 1-PHOSPHOTRAN |
| 14: | 9dqm-B | 62.2 | 0.6 | 421 | 421 | 100 | PDB | MOLECULE: PYROPHOSPHATE--FRUCTOSE 6-PHOSPHATE 1-PHOSPHOTRAN |
| 15: | 9mf6-D | 62.2 | 0.5 | 417 | 417 | 100 | PDB | MOLECULE: PYROPHOSPHATE--FRUCTOSE 6-PHOSPHATE 1-PHOSPHOTRAN |
| 16: | 9mf6-C | 62.1 | 0.6 | 420 | 420 | 100 | PDB | MOLECULE: PYROPHOSPHATE--FRUCTOSE 6-PHOSPHATE 1-PHOSPHOTRAN |
| 17: | 9dqm-A | 61.9 | 0.7 | 420 | 420 | 100 | PDB | MOLECULE: PYROPHOSPHATE--FRUCTOSE 6-PHOSPHATE 1-PHOSPHOTRAN |
| 18: | 9med-A | 61.6 | 0.8 | 417 | 417 | 100 | PDB | MOLECULE: PYROPHOSPHATE--FRUCTOSE 6-PHOSPHATE 1-PHOSPHOTRAN |
| 19: | 9mec-A | 61.6 | 0.9 | 422 | 422 | 100 | PDB | MOLECULE: PYROPHOSPHATE--FRUCTOSE 6-PHOSPHATE 1-PHOSPHOTRAN |
| 20: | 9mdt-A | 61.5 | 0.8 | 417 | 417 | 100 | PDB | MOLECULE: PYROPHOSPHATE--FRUCTOSE 6-PHOSPHATE 1-PHOSPHOTRAN |
| 21: | 9mea-A | 61.3 | 0.8 | 418 | 418 | 100 | PDB | MOLECULE: PYROPHOSPHATE--FRUCTOSE 6-PHOSPHATE 1-PHOSPHOTRAN |
| 22: | 9mf6-A | 60.3 | 1.0 | 418 | 418 | 100 | PDB | MOLECULE: PYROPHOSPHATE--FRUCTOSE 6-PHOSPHATE 1-PHOSPHOTRAN |
| 23: | 9cit-D | 38.2 | 2.7 | 357 | 378 | 31 | PDB | MOLECULE: 6-PHOSPHOFRUCTOKINASE; |
| 24: | 9cis-C | 37.9 | 2.6 | 355 | 377 | 30 | PDB | MOLECULE: 6-PHOSPHOFRUCTOKINASE; |
| 25: | 9cir-D | 37.8 | 2.7 | 352 | 372 | 31 | PDB | MOLECULE: 6-PHOSPHOFRUCTOKINASE; |
| 26: | 9cir-C | 37.8 | 2.7 | 353 | 377 | 31 | PDB | MOLECULE: 6-PHOSPHOFRUCTOKINASE; |
| 27: | 9cir-A | 37.3 | 2.7 | 354 | 385 | 31 | PDB | MOLECULE: 6-PHOSPHOFRUCTOKINASE; |
| 28: | 9cit-C | 37.3 | 2.7 | 352 | 380 | 31 | PDB | MOLECULE: 6-PHOSPHOFRUCTOKINASE; |
| 29: | 9cis-B | 37.2 | 2.7 | 351 | 375 | 31 | PDB | MOLECULE: 6-PHOSPHOFRUCTOKINASE; |
| 30: | 9cis-A | 37.2 | 2.9 | 361 | 384 | 30 | PDB | MOLECULE: 6-PHOSPHOFRUCTOKINASE; |
| 31: | 9cit-B | 37.1 | 2.8 | 357 | 385 | 31 | PDB | MOLECULE: 6-PHOSPHOFRUCTOKINASE; |
| 32: | 9cir-B | 37.0 | 3.0 | 362 | 391 | 31 | PDB | MOLECULE: 6-PHOSPHOFRUCTOKINASE; |
| 33: | 9cis-D | 36.7 | 2.6 | 345 | 366 | 31 | PDB | MOLECULE: 6-PHOSPHOFRUCTOKINASE; |
| 34: | 6qu5-F | 36.7 | 2.6 | 346 | 456 | 29 | PDB | MOLECULE: ATP-DEPENDENT 6-PHOSPHOFRUCTOKINASE; |
| 35: | 9cit-A | 36.5 | 2.8 | 358 | 389 | 30 | PDB | MOLECULE: 6-PHOSPHOFRUCTOKINASE; |
| 36: | 5xz7-A | 36.3 | 2.0 | 318 | 322 | 25 | PDB | MOLECULE: ATP-DEPENDENT 6-PHOSPHOFRUCTOKINASE; |
| 37: | 3pfk-A | 36.0 | 2.0 | 317 | 319 | 27 | PDB | MOLECULE: PHOSPHOFRUCTOKINASE; |
| 38: | 5xz8-A | 35.9 | 1.9 | 314 | 317 | 26 | PDB | MOLECULE: ATP-DEPENDENT 6-PHOSPHOFRUCTOKINASE; |
| 39: | 4a3s-B | 35.8 | 1.9 | 316 | 319 | 25 | PDB | MOLECULE: 6-PHOSPHOFRUCTOKINASE; |
| 40: | 1pfk-B | 35.8 | 1.9 | 316 | 320 | 28 | PDB | MOLECULE: PHOSPHOFRUCTOKINASE; |
| 41: | 4a3s-A | 35.8 | 1.9 | 316 | 319 | 25 | PDB | MOLECULE: 6-PHOSPHOFRUCTOKINASE; |
| 42: | 6qu3-D | 35.8 | 2.5 | 351 | 461 | 28 | PDB | MOLECULE: ATP-DEPENDENT 6-PHOSPHOFRUCTOKINASE; |
